# Mapping the genetic landscape of iron metabolism uncovers the SETD2 methyltransferase as a modulator of iron flux

**DOI:** 10.1101/2025.02.26.640314

**Authors:** Anthony W. Martinelli, Chun-Pei Wu, Tristan Vornbäumen, Niek Wit, Jia J. Sia, Anneliese O. Speak, James A. Nathan

## Abstract

Cellular iron levels must be tightly regulated to ensure sufficient iron for essential enzymatic functions whilst avoiding the harmful generation of toxic species. Here, to better understand how iron levels are controlled, we carry out genome-wide mutagenesis screens in human cells. Alongside mapping known components of iron-sensing, we determine the relative contributions of iron uptake, iron recycling, ferritin breakdown, and mitochondrial flux in controlling the labile iron pool. We also identify SETD2, a histone methyltransferase, as a chromatin modifying enzyme that controls intracellular iron availability through ferritin breakdown. Functionally, we show that SETD2 inhibition or cancer-associated SETD2 mutations render cells iron deficient, thereby driving resistance to ferroptosis, and potentially explaining how some tumours evade anti-tumoural immunity.

## Introduction

Strict regulation of cellular iron levels is essential. Sufficient iron must be present within cells to ensure that it can be incorporated into life-critical proteins and facilitate redox reactions, whilst avoiding the generation of harmful reactive oxygen species. Cells must therefore be able to sense and respond rapidly to changes in iron abundance.

Iron-sensing is principally mediated at a post-transcriptional level by the iron regulatory proteins 1 and 2 (IRP1 and IRP2) (*1–5*). In conditions of iron depletion, IRPs bind to iron responsive elements (IREs) within the 5-prime (5’) or 3-prime (3’) regions of gene transcripts involved in iron metabolism. Binding within the 5’ region inhibits translation, whereas 3’ IREs promote mRNA stabilization and increased translation (*6–11*). The co-ordination of this IRP response maintains iron homeostasis by regulating pathways critical for iron uptake, recycling, and storage (*12*).

Most cellular iron is taken up in a transferrin-bound ferric (Fe^3+^) form via the transferrin receptor (TfR) (*13*). Iron is then converted to a ferrous (Fe^2+^) form in early endosomes before incorporation into proteins or delivery to the mitochondria (*14–16*). Excess iron is either exported from the cell, via ferroportin, or is stored bound to ferritin – a protective cage that mitigates against the risks of this reactive metal (*17–20*). Reactive iron can be released from ferritin via a specialized form of autophagy, termed ferritinophagy, which requires a specific cargo receptor, NCOA4 (*21*).

Aside from the essential role of iron in sustaining cellular redox reactions, work over the last decade has identified that iron is necessary for a specialized form of cell death, ferroptosis, which is characterized by a lipid peroxidation cascade (*22–25*). Inducing ferroptosis provides a promising therapeutic approach to treat cancers and it is also noteworthy that ferroptosis can be induced by CD8+ T cells as an anti-tumour mechanism (*26–29*). Furthermore, induction of ferroptosis in cancer cells upregulates the expression of ULBPs which are activating ligands for natural killer (NK) cells via the NKG2D receptor (*30*). Thus, there is potential for a synergistic effect both directly and indirectly via activation of innate and adaptive cytotoxic immune cells.

While the relationship between iron uptake and ferritin breakdown is established, many aspects of cellular iron metabolism remain unknown, including the exact nature of free iron within the cell (the labile iron pool, LIP), how iron is trafficked and incorporated into proteins, and what role, if any, the nucleus plays in coordination of these processes (*12*).

Additionally, how modulating cellular iron impacts ferroptosis and whether the dysregulation of iron homeostasis that occurs in cancers protects against ferroptosis is unclear (*31–33*).

Here, we establish sensitive and dynamic reporters for measuring cellular iron flux and use these tools to identify the key genes and pathways which regulate iron metabolism. Alongside mapping the landscape of cellular iron homeostasis, we uncover the histone methyltransferase SETD2 as a novel transcriptional regular of cellular iron flux. We also show that SETD2 mutations, which occur frequently in renal cell carcinomas, render these cancer cells resistant to ferroptosis, thereby uncovering a pro-oncogenic mechanism dependent on intracellular iron regulation.

## Results

### IRP2-Clover reporter cells provide a sensitive and dynamic readout for intracellular iron levels

To map the landscape of intracellular iron metabolism we first generated tools to measure iron flux. We focused on IRP2 as it provides the main cytosolic sensing arm of intracellular iron availability and IRP2 protein levels directly correspond to iron deficiency (*3*). In conditions of iron repletion, IRP2 is ubiquitinated and targeted for proteasomal degradation by the FBXL5 Skp1-Cullin-F-box ubiquitin ligase complex in an iron-dependent manner (**Fig. 1A**) (*34, 35*). Conversely in conditions of iron depletion, IRP2 is stabilized and accumulates, binding iron responsive elements (IREs) in the 5’ or 3’-UTRs of relevant mRNAs to exert post-transcriptional control of iron-relevant genes (*3*).

**Fig. 1:**
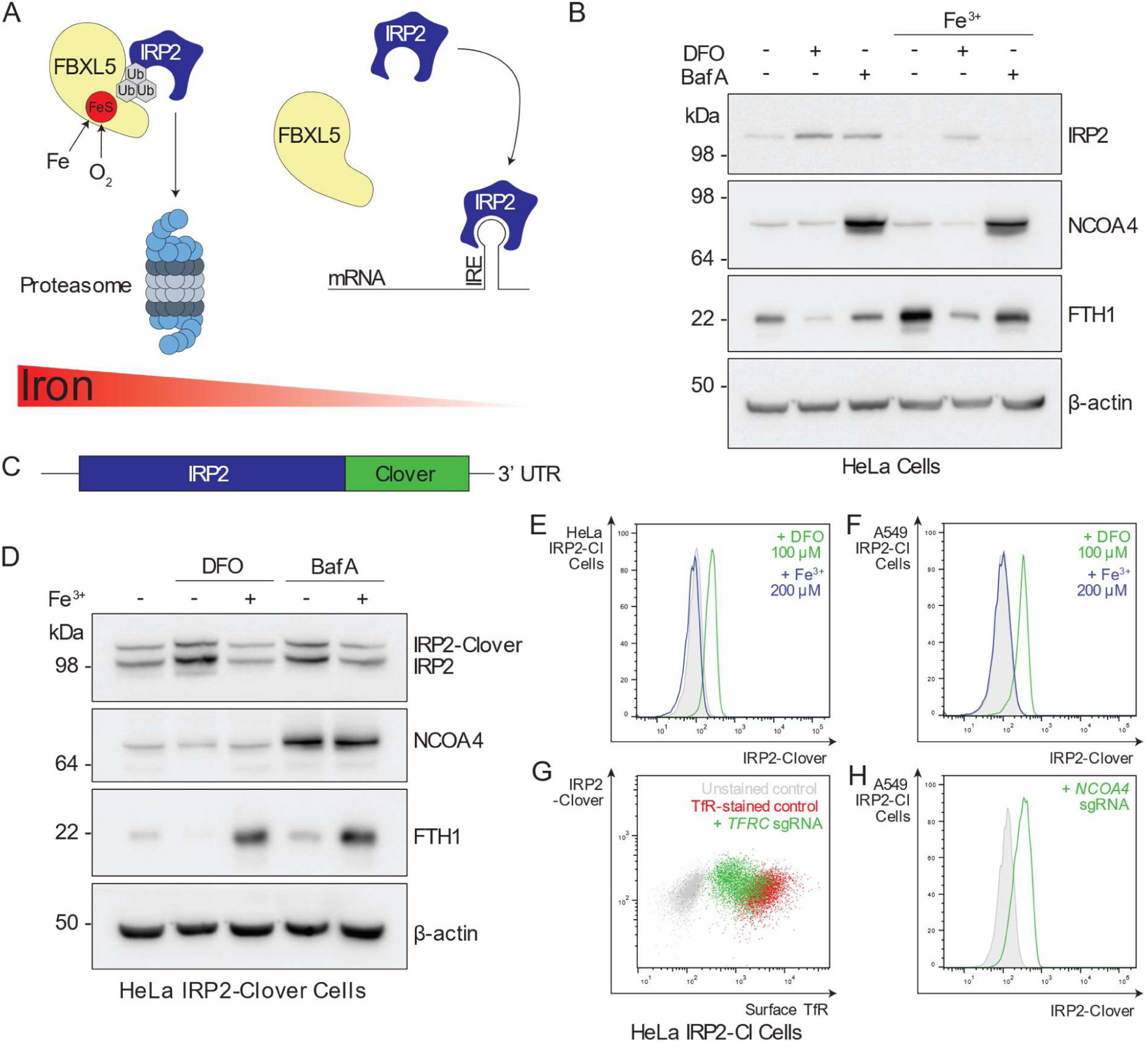
IRP2-Clover reporter cells provide a sensitive and dynamic readout for intracellular iron levels. (**A**) In conditions of iron repletion, IRP2 is targeted for proteasomal degradation by FBXL5 in an oxygen-dependent manner, whereas when iron is deplete, IRP2 binds iron-responsive elements to exert post-transcriptional control of cellular iron homeostasis. (**B**) HeLa cells were treated with DFO (100 μM, 24 hr) or BafA (10 nM, 24 hr) with or without ferric iron supplementation (FAC, 100 μM, 24 hr) and protein levels of IRP2, ferritin heavy chain (FTH1), ferritinophagy cargo receptor NCOA4 and β-actin assessed by immunoblot (n=3). (**C**) Schematic of the IRP2-Clover reporter construct. (**D**) HeLa IRP2-Clover cells were treated with iron depletion by BafA (10 nM, 20 hr) or DFO (100 μM, 20 hr) with or without addition of iron. Cells were analysed by immunoblot for IRP2, NCOA4, FTH1 and β-actin (n=4). (**E**) HeLa IRP2-Clover cells were treated with iron depletion by DFO (200 μM, 20 hr) with or without addition of iron and analysed by flow cytometry (n=5). (**F**) A549 IRP2-Clover cells were treated with iron depletion by DFO (200 μM, 20 hr) with or without addition of iron and analysed by flow cytometry (n=3). (**G**) HeLa IRP2-Clover cells were transduced with sgRNA targeting *TFRC* with cell surface antibody staining for TfR and analysis by flow cytometry (n=3). (**H**) Cells were transduced with sgRNA targeting *NCOA4* before analysis by flow cytometry (n=3). *IRP2-Clover, IRP2-Cl*.

We used two strategies to determine the utility of IRP2 as a readout for intracellular iron availability. We first determined if iron availability could be measured by IRP2 intracellular antibody staining and flow cytometry. HeLa cells treated with the iron-chelator desferrioxamine (DFO) (100 µM, 24 hr) showed an increase in IRP2 levels by both immunoblot and flow cytometry (**Fig. 1B** and **fig. S1A**). IRP2 accumulation was reversed by addition of exogenous ferric iron (ferric ammonium citrate, FAC, 100 µM, 24 hr). Additionally, inhibition of iron flux using the v-ATPase inhibitor, bafilomycin (BafA, 10 nM, 24 hr) resulted in IRP2 accumulation which was reversible by addition of iron, confirming the utility of IRP2 as a robust readout of iron levels (**Fig. 1B** and **fig. S1B**).

Secondly, we developed HeLa and A549 clonal knock-in IRP2-Clover cell lines by endogenously tagging the C-terminus of IRP2 with the fluorescent protein Clover (**Fig. 1C**; **fig. S1C**). Immunoblot and flow cytometry confirmed that IRP2-Clover accumulation occurred following iron depletion (iron chelation or v-ATPase inhibition) (**Fig. 1D-F** and **fig. S1D and E**) (*36, 37*), and ferric iron supplementation (FAC, 200 µM, 24 hr) reversed IRP2-Clover accumulation (**Fig. 1D-F** and **fig. S1D and E**). To determine if the IRP2-Clover cells would also respond to genetic manipulation of genes involved in iron regulation, we knocked out *TFRC* and *NCOA4* using sgRNA and observed that IRP2-Clover levels increased when either TfR uptake or ferritinophagy were prevented (**Fig. 1G and H** and **fig. S1F**).

### Genome-wide mutagenesis screens define the key regulators of intracellular iron metabolism

Having established the utility of IRP2 as a sensitive readout for cellular iron regulation, we used CRISPR/Cas9 mutagenesis screens to find the key genes and pathways regulating iron homeostasis. A549 or HeLa IRP2-Clover cells, stably expressing Cas9, were transduced with genome-wide sgRNA knockout (KO) libraries and underwent iterative fluorescence activated cell sorting (FACS) to enrich for IRP2-Clover^HIGH^ levels, thus identifying genes which, when lost, promote IRP2 accumulation and by implication lead to reduced iron availability (**Fig. 2A**). Two sgRNA libraries were used to mitigate against differences in sgRNA design: the Whitehead library was transduced into HeLa IRP2-Clover cells, whereas the TKOv3 library was used in the A549 IRP2-Clover cells (*38–40*). We also compared the findings of the knock-in IRP2 reporters with an intracellular FACS screen, using antibody staining of endogenous IRP2 in HeLa cells (Whitehead or TKOv1 library) (*38–40*) (**fig. S2A**). For all screens, DNA was extracted at several time points post mutagenesis (day 7-8, “early” and day 14-17, “late”) to maximize the sensitivity of our approach and prevent the loss of genetic mutations that have more severe phenotypes (e.g. decreased cell growth or death).

**Fig. 2:**
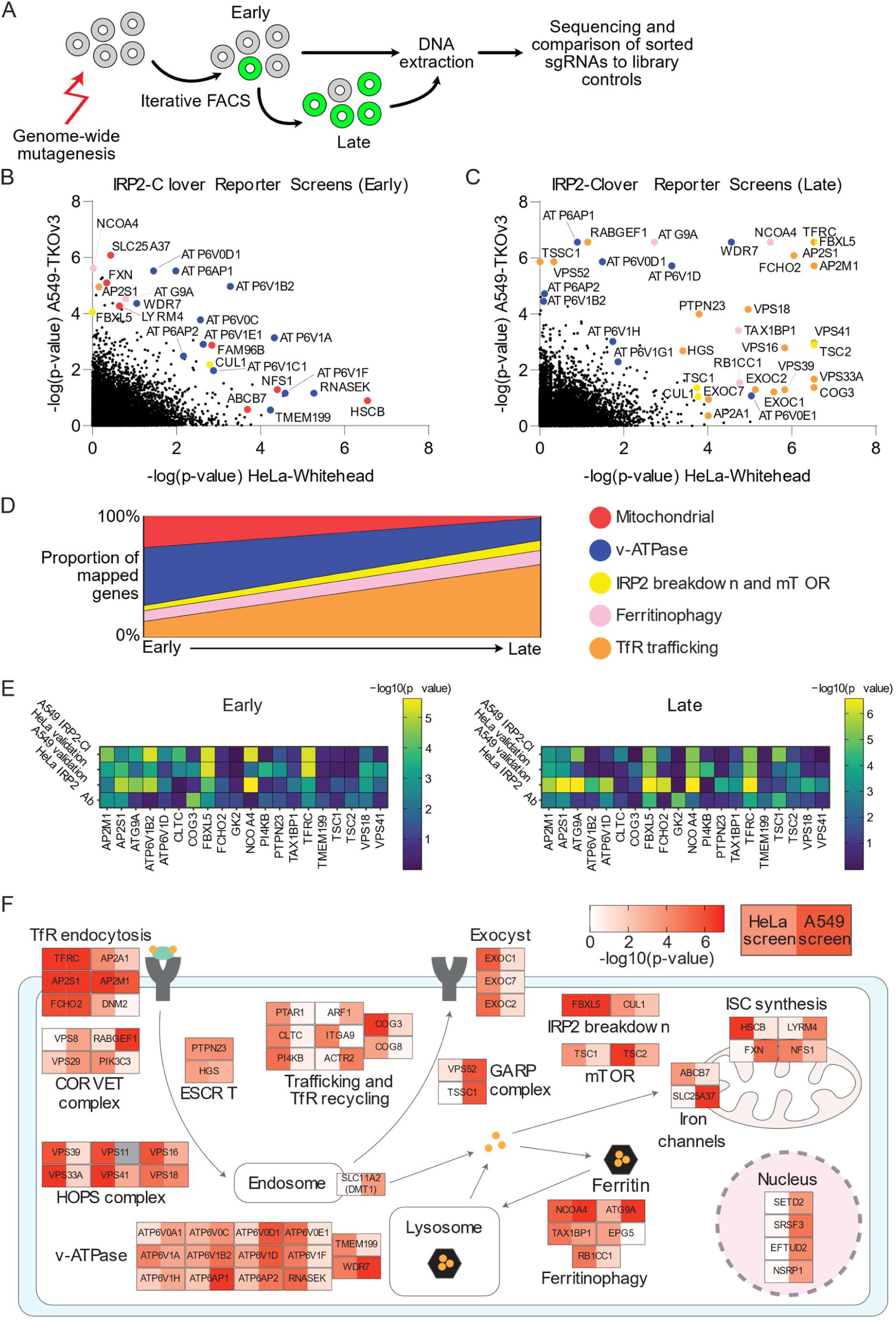
Genome-wide mutagenesis screens define the key regulators of intracellular iron metabolism. (**A**) HeLa or A549 IRP2-Clover cells expressing Cas9 were mutagenised with genome-wide sgRNA libraries, selected for lentiviral integration and underwent FACS for Clover^HIGH^ cells after 8 days. Sorted cells were split between lysis for immediate DNA extraction and expansion for a second sort (day 16-18). DNA from phenotypically non-selected library cells was extracted for comparison and cells were pooled prior to any selection event. (**B**, **C**) Bubble plots showing screen hits (genes overrepresented in sorted cells, as calculated by MAGeCK) identified in the A549 IRP2-Clover-TKOv3 screen (y-axis) compared to the HeLa IRP2-Clover-Whitehead screen (x-axis) at early (**B**, day 8) and late (**C**, day 16 or day 18 respectively) time points. Genes with related functions are highlighted. (**D**) Genes identified in either screen as top hits (-log(p-value)>3) at early or late time points were manually annotated for function to assess changes over time, with genes of unknown function excluded (49/72 unknown at early time point and 44/104 unknown at late time point). (**E**) Heat maps illustrating relative -log(p-value) selected hits across primary screens in HeLa (antibody screen, TKOv1) and A549 IRP2-Clover (TKOv3) and secondary validation sub-pooled library screens in HeLa and A549 IRP2-Clover cells. (**F**) Across the HeLa IRP2-Clover-Whitehead and A549 IRP2-Clover-TKOv3 screens (both early and late time points) 150 unique genes were identified as registering a -log(p-value) >3 by MAGeCK analysis. Of these, 61 mapped to complexes and pathways with identifiable roles in cellular iron metabolism. For each gene the colour in the left box represents the highest -log(p-value) of the two time points assessed for the HeLa IRP2-Clover screen and the colour in the right box corresponds to the equivalent value for the A549 IRP2-Clover screen. *IRP2-Cl=IRP2-Clover*.

All mutagenesis screens identified key positive controls, including *FBXL5* and *TFRC* (**Fig. 2B** and **fig. S2B and C**), validating the approach. In addition, we identified a conserved group of genes potentially regulating iron levels (**Fig. 2B and C** and **fig. S2B and C**). The IRP2-Clover reporter screens were more sensitive for detecting candidate genes compared to the antibody screens. In addition, the timing of flow cytometry sort post-mutagenesis influenced the genes identified. For instance, genes annotated as mitochondrial regulators were found following the first round of FACS, whilst genes relating to endocytosis of the transferrin receptor were proportionally higher at later time points (**Fig. 2D**), likely reflecting the toxicity of perturbing mitochondrial function on cell growth.

We next generated a sub-pooled sgRNA library of the top hits identified in the screens (targeting 184 genes) to validate our findings in both HeLa and A549 IRP2-Clover cells (**fig. S2D** and **data S1**). By comparing the top hits across the primary and secondary screens we identified a core set of genes that were reproducibly enriched for sgRNA (**Fig. 2D**). Whilst key regulatory genes (e.g. *FBXL5, NCOA4* and *TFRC*) could be identified in all screens, other genes were identified at different time points post-mutagenesis, and in different cell lines, consistent with the temporal effects previously observed (**Fig. 2B-E**) and suggested some cell-type specific responses.

Lastly, we mapped the findings of the screens using gene ontology and known function. Gene ontology analysis confirmed the significant pathways identified by manual annotation including ‘synaptic vesicle cycle’ as the most represented biological processes, ‘endosome membrane’ (containing the v-ATPase) as the most prominent cellular component, and ‘iron uptake and transport’ as the top Reactome pathway (**fig. S2E**) (*41–43*). Of the top 150 genes identified (-log(p-value) >3), nearly half (65 genes) could be mapped by function and grouped into key pathways of cellular iron metabolism (**Fig. 2F**). Transferrin receptor internalization and recycling dominated the top hits, with additional components such as the HOPS (homotypic fusion and protein sorting) and GARP (Golgi-associated retrograde protein) complexes identified relating to transferrin recycling (*44, 45*). The involvement of the v-ATPase in reducing ferric to ferrous iron was also apparent, as was the requirement for ferritinophagy (*21, 36*). Less studied areas that were identified were the regulation of iron between the cytosol and mitochondria, the involvement of the exocyst complex, and the role of several nuclear localizing genes, including *SETD2* (**Fig. 2F**). Therefore, we chose to explore the relative contribution of the exocyst complex and mitochondria to iron metabolism, and the role of SETD2.

### The exocyst complex and mitochondrial iron metabolism influence cellular iron flux

The exocyst is involved in trafficking of vesicles to the plasma membrane for secretion (*46*). Three genes encoding subunits of the exocyst (*EXOC1, EXOC2*, and *EXOC7*) were identified in our screens (**Fig. 2B and F**), but the role of this complex in intracellular iron regulation was not clear. The exocyst may be required for TfR recycling in erythroblasts, but its role outside of erythropoiesis, where iron requirements are particularly high, has not been described (*47*). Therefore, we chose to knock down *EXOC1* by lentiviral shRNA transduction and measured the IRP2 response in A549 IRP2-Clover cells.

Both endogenous IRP2 and IRP2-Clover levels increased following EXOC1 depletion (**Fig. 3A**). Similar findings were identified using shRNA targeting *EXOC1* in A549 IRP2-Clover reporter cells (**Fig. 3B**). However, while increased IRP2 levels suggested perturbation of iron flux when the exocyst was depleted, we did not observe any change in steady state levels of cell surface TfR after knockout of *EXOC1* in either A549 or HeLa IRP2-Clover cells (**Fig. 3C**). Therefore, we used a transferrin (Tf) uptake and recycling assay with fluorescently-labelled Tf (Tf-AF647) to establish how the exocyst may be implicated in intracellular iron regulation (**Fig. 3D**). HeLa cells were pulsed with Tf-AF647 for 5 min and labelled-Tf and cell-surface TfR measured after 30 or 60 min. Tf-AF647 levels reduced at 30 and 60 min in control HeLa cells, consistent with rapid recycling of Tf to the plasma membrane (**Fig. 3D**). However, high levels of Tf-A647 were retained in EXOC1 depleted cells, indicating that TfR recycling was impaired (**Fig. 3D**). TfR levels also transiently increased at the cell surface in pulsed cells which had EXOC1 loss (**Fig. 3D**). Therefore, loss of the exocyst complex results in impaired iron flux due to defective TfR recycling, even in cells without enhanced iron requirements.

**Fig. 3:**
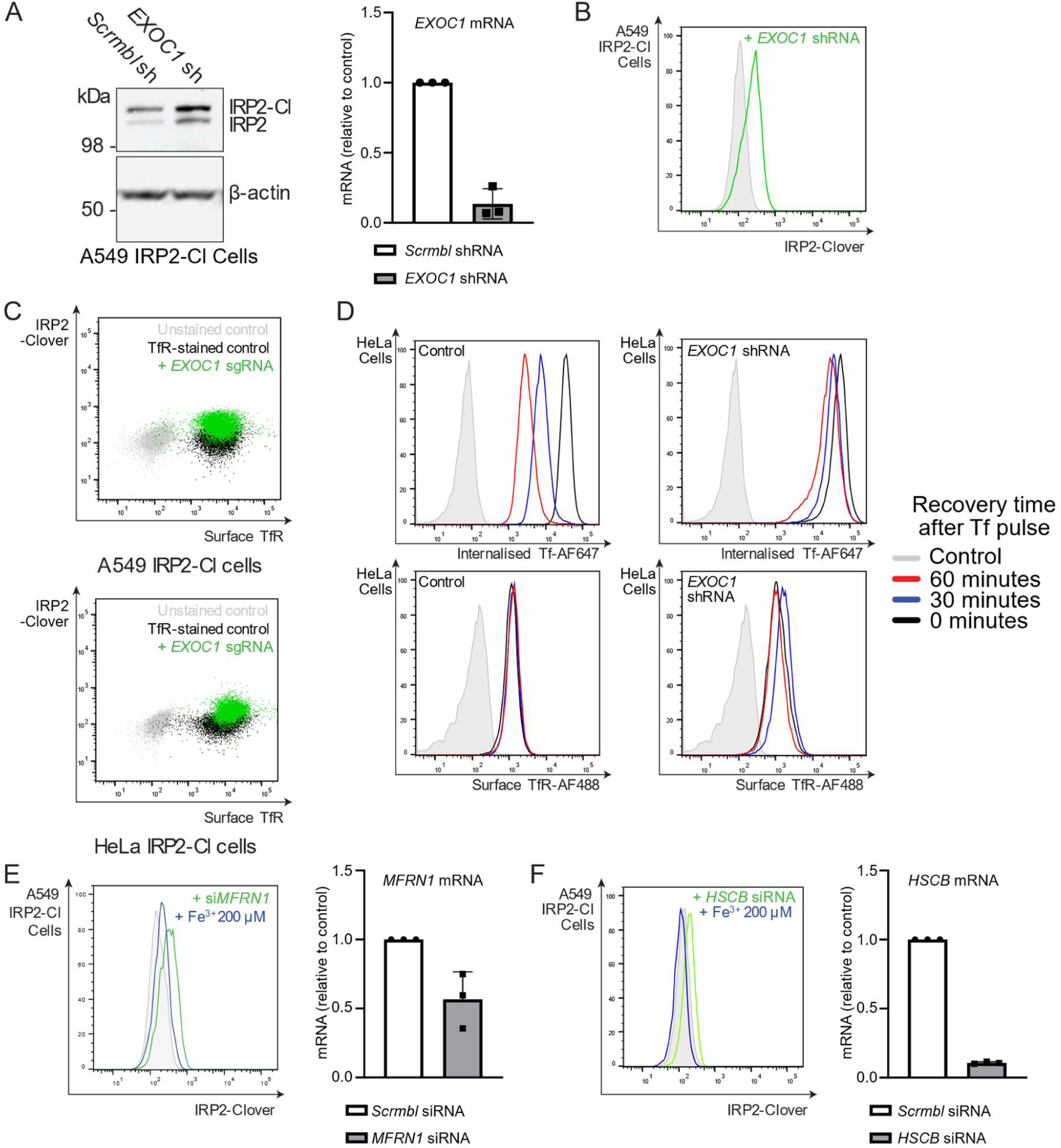
The exocyst complex and mitochondrial iron metabolism influence cellular iron flux. (**A**) A549 IRP2-Clover cells were transduced with shRNA targeting *EXOC1* and underwent immunoblot for IRP2 and β-actin. *EXOC1* depletion was confirmed by RT-qPCR (n=3). (**B**) A549 IRP2-Clover cells were transduced with shRNA targeting *EXOC1* and underwent flow cytometry for measurement of IRP2-Clover levels (n=3). (**C**) A549 IRP2-Clover and HeLa IRP2-Clover cells were transduced with sgRNA target *EXOC1* and stained with antibody targeting surface TfR before analysis by flow cytometry (n=2 for each cell line). (**D**) TfR uptake and recycling. HeLa cells with or without knockdown of *EXOC1* by shRNA were serum-starved for 45 min, incubated with 5 µg/ml Tf-AF647 for 5 min, washed with PBS and recovered in serum free medium for the indicated time periods. Internalized Tf-AF647 was assessed by flow cytometry. TfR surface levels were detected by flow cytometry (n=3, note controls identical for both conditions). (**E**) A549 IRP2-Clover cells were transfected with siRNA targeting *MFRN1* ± ferric iron (FAC 200 µM, 24 hr) before analysis by flow cytometry (left panel). Knockdown was confirmed by RT-qPCR (right panel) (n=3). (**F**) A549 IRP2-Clover cells were transfected twice with siRNA targeting *HSCB* (48 hr and 24 hr prior to analysis by flow cytometry) ± ferric iron (FAC 200 µM, 24 hr). Knockdown was confirmed by RT-qPCR (right panel) (n=3). *IRP2-Clover, IRP2-Cl*.

The identification of genes involved in mitochondrial iron metabolism suggested that the mitochondria themselves can impact cytosolic iron flux. Related genes identified in our screens included the mitochondrial iron sulfur cluster (ISC) chaperone, *HSCB*, and the mitochondrial iron importer *SLC25A37* (Mitoferrin-1, *MFRN1*). We depleted HSCB and Mitoferrin-1 using siRNA and confirmed an increase in IRP2-Clover levels (**Fig. 3E, F**). Supplementing the media with excess iron restored IRP2-Clover levels, confirming that normal intracellular iron flux was impaired. Therefore, both impaired ISC synthesis or reduced mitochondrial iron levels can drive accumulation of IRP2, potentially driving increased cytosolic iron levels as a compensatory response.

### Loss of the histone methylase SETD2 results in an iron-dependent IRP2 response

Cellular iron homeostasis is not known to be regulated by transcription. Therefore, the identification of *SETD2,* a methyltransferase responsible for adding the H3 lysine 36 trimethylation (H3K36me3) mark (*48, 49*), as a hit within the genetic screens was intriguing (**Fig. 4A** and **fig. S3A**). SETD2 also associates with the spliceosome (*50*), and several spliceosome genes were identified in our IRP2-Clover reporter screens (*SRSF3, NSRP1, EFTUD2*, and *HNRNPU*), suggesting a possible transcriptional role of a SETD2 complex in iron metabolism (**Fig. 4A and B**). Therefore, we examined if SETD2 loss altered iron flux and if this was dependent on its catalytic activity.

**Fig. 4:**
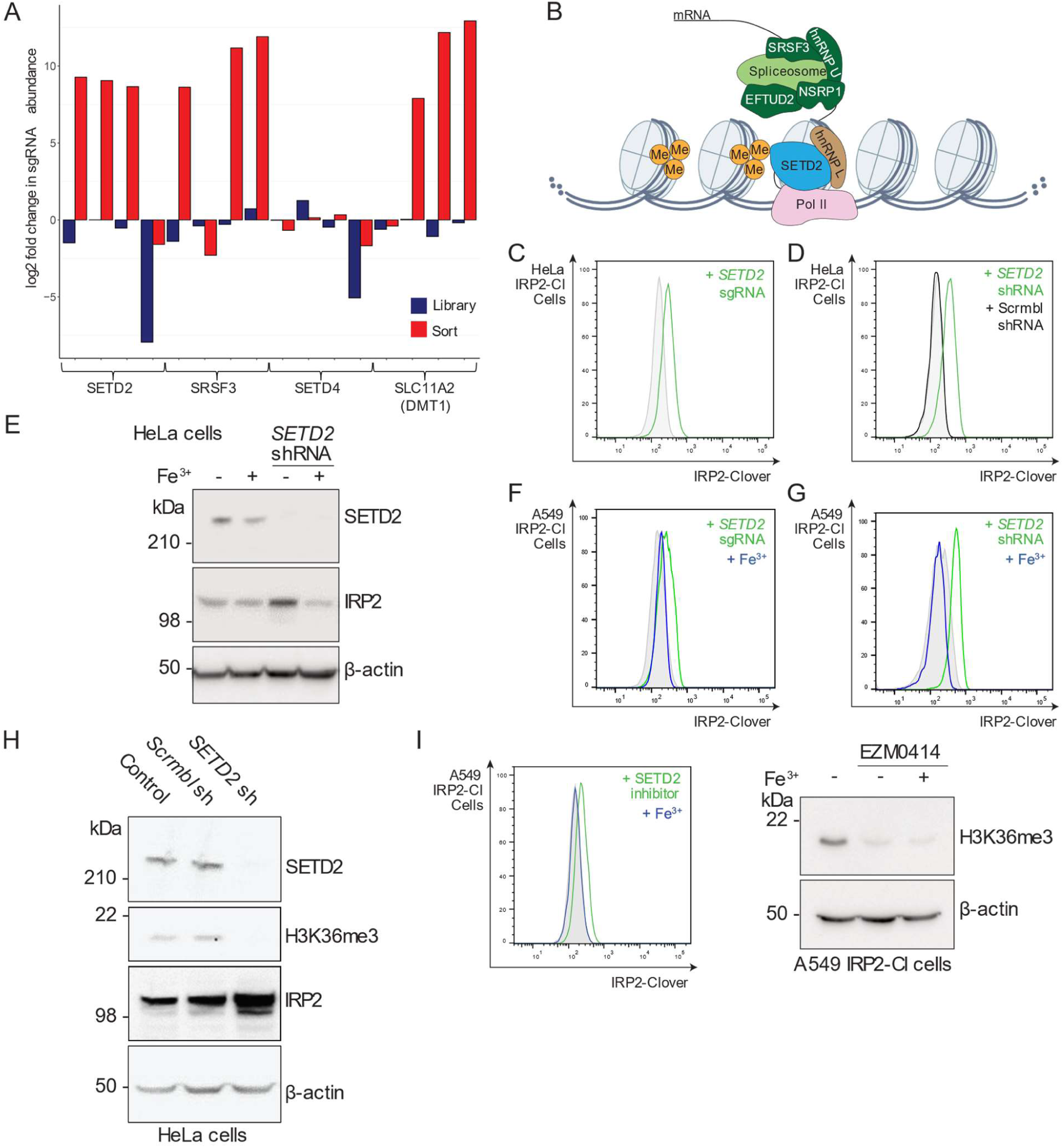
SETD2-mediated IRP2 accumulation is dependent on enzymatic activity. (**A**) Bar chart detailing the log2-fold change in individual sgRNA counts for *SETD2* in the A549 IRP2-Clover CRISPR screen, as well as an associated splicing regulator (*SRSF3*). *DMT1* was included as a positive control, and *SETD4* as a negative control. (**B**) Schematic illustrating the canonical role of SETD2 in H3K36me3 and regulation of splicing. Spliceosome components enriched for sgRNA in the screens are shown in dark green. (**C**, **D**) HeLa IRP2-Clover cells were transduced with sgRNA targeting *SETD2* (**C**) (n=4), or shRNA targeting *SETD2* with a shRNA scrambled control (**D**) (representative of at least three biological replicates). IRP2-Clover fluorescence measured by flow cytometry. (**E**) HeLa cells were transduced with shRNA targeting *SETD2* and treated with or without ferric iron supplementation (FAC, 200 μM) for 24 hr prior to analysis by immunoblot (n=4). (**F**, **G**) A549 IRP2-Clover cells were transduced with sgRNA (n=3) or shRNA (n=4) targeting *SETD2* and treated with or without ferric iron supplementation (FAC, 200 μM, 24 hr) prior to analysis by flow cytometry. (**H**) HeLa cells were transduced with shRNA targeting SETD2 and cells were lysed before analysis by immunoblot for levels of SETD2, H3K36me3, IRP2, and β-actin (n=4). (**I**) A549 IRP2-Clover cells were treated with EZM0414 (200 nM, 48 hr), with or without ferric iron supplementation (FAC 200 µM, 24 hr) before analysis by flow cytometry for IRP2-Clover levels (left panel) and immunoblot for H3K36me3 and β-actin (right panel) (n=3). *IRP2-Cl=IRP2-Clover*.

We first validated that SETD2 depletion increased IRP2 levels using sgRNA or shRNA-mediated depletion in IRP2-Clover reporter cells (**Fig. 4C and D**). A similar increase in untagged endogenous IRP2 following SETD2 depletion was seen in HeLa cells (**Fig. 4E**). We also observed IRP2 accumulation in HAP1, 786-O and MCF7 cells lines when SETD2 was depleted, confirming the generalizability of our findings (**fig. S3B-D**). IRP2 accumulation following SETD2 loss was dependent on iron flux, as ferric iron (FAC, 200 µM, 24 hr) reduced IRP2 levels in both HeLa cells and A549 IRP2-Clover reporter cells (**Fig. 4E-G**). Therefore, depletion of SETD2 results in accumulation of IRP2, in an iron-responsive manner.

As the main function of SETD2 relates to its methyltransferase activity (*48, 49*), we examined if SETD2 inhibition altered iron flux. SETD2 shRNA-mediated depletion or inhibition with EZM-0414 (200 nM, 48 hr) (*51*) decreased H3K36me3 levels (**Fig. 4H** and **I**), as expected. Moreover, IRP2-Clover levels increased following SETD2 inhibition in an iron-dependent manner, similarly to SETD2 depletion (**Fig. 4I**). Therefore, the SETD2 enzymatic activity is required for IRP2 regulation.

While the canonical role of SETD2 relates to H3K36me3 (*48, 49*), non-histone methylation by SETD2 has been reported on microtubules and actin, where it alters polymerization (*52, 53*). However, the subcellular localization of SETD2 argued against a non-canonical role of SETD2, as both immunofluorescence and subcellular fractionation assays showed SETD2 was bound primarily to chromatin in HeLa cells (**fig. S4A and B**). In addition, we did not observe IRP2 accumulation with inhibition of microtubule polymerization (CK-869, 200 nM, 24 hr) or reversal of IRP2 levels with the actin polymerization promoter Jasplakinolide (100 nM, 24 hr) (**fig. S4C**). Together, these findings support SETD2 acting via its canonical H3K36me3 function to drive IRP2 accumulation.

### SETD2 deficiency results in defective ferritinophagy

To understand how SETD2 may influence iron flux, we examined which aspect of intracellular iron metabolism was impaired when SETD2 was depleted. While total cell iron levels remained constant (**Fig. 5A**), a striking finding was the accumulation of the ferritinophagy cargo receptor, NCOA4, in SETD2 deficient cells (**Fig. 5B**). Perturbations in ferritinophagy may arise as a compensatory mechanism for impaired iron availability (*54–58*), and as SETD2 modulates transcription, we first considered whether the TfR pathway may be impaired following SETD2 loss. However, *TFRC* mRNA levels were not altered following SETD2 depletion (**Fig. 5C** and **fig. S5A**). Moreover, cell-surface levels of TfR levels remained similar to those seen in SETD2-replete HeLa or A549 IRP2-Clover cells (**Fig. 5C** and **fig. S5B**). Tf uptake and recycling was also not altered by SETD2 loss (**Fig. 5D**). Therefore, as the Tf pathway could not account for the alterations in iron flux following SETD2 depletion, we focused on ferritin breakdown or ferritinophagy.

**Fig. 5:**
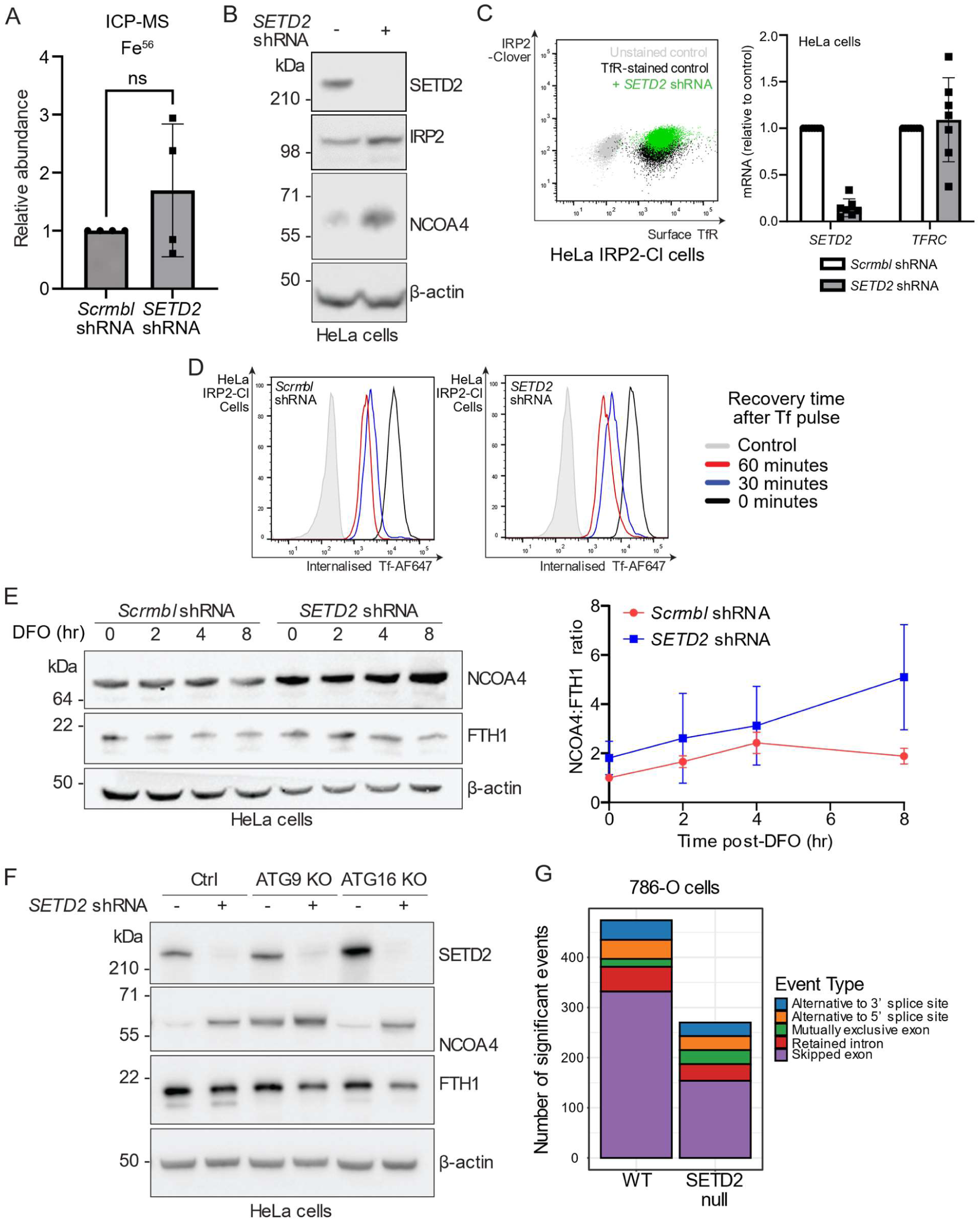
Loss of SETD2 results in increased NCOA4 and altered ferritinophagy. (**A**) HeLa cells were transduced with shRNA targeting *SETD2* or a scrambled control. Cells were lysed in nitric acid before analysis by ICP-MS. Duplicate cell cultures were counted and measurements normalized to cell number (n=4 in technical triplicate; p=0.31, paired t test). (**B**) HeLa cells were transduced with shRNA targeting *SETD2* and protein levels of SETD2, IRP2, NCOA4 and β-actin measured by immunoblot (representative of at least three biological replicates). (**C**) HeLa and HeLa IRP2-Clover cells were transduced with shRNA targeting *SETD2* and underwent flow cytometry for surface TfR and IRP2-Clover (n=3). Transcript levels of *SETD2* and *TFRC* were measured by RT-qPCR (n=7). (**D**) TfR uptake and recycling. SETD2 was depleted in HeLa IRP2-Clover cells using shRNA, along with a shRNA scrambled control. Cells were serum-starved for 5 min, incubated with Tf-AF647 and recovered in serum free medium for the indicated time points. Internalized Tf-AF647 was assessed by flow cytometry (n=3). (**E**) Control or SETD2 depleted HeLa cells (shRNA) were treated with DFO (100 μM) for 0-8 hr and levels of NCOA4, FTH1 and β-actin measured by immunoblot (n=3). Immunoblots were quantified and normalized to loading and 0 hr time points before calculation of the NCOA4:FTH1 ratio. (**F**) HeLa null ATG9, ATG16, or paired controls were transduced with shRNA targeting *SETD2* and immunoblot performed for SETD2, NCOA4, FTH1 and β-actin (representative of at least three biological replicates). (**G**) Publicly available RNA-seq data from wild type and *SETD2* stable knockout (SETD2 null) 786-O cells (GSE150609) was analysed using rMATS, and splicing events quantified. *IRP2-Cl=IRP2-Clover*.

NCOA4 accumulation is typically only observed when ferritinophagy is impaired, as NCOA4 is co-degraded with ferritin within the lysosome (21, 36). Therefore, to assess if SETD2 loss impaired ferritinophagy, we measured the kinetics of NCOA4 and ferritin (ferritin heavy chain, FTH1) levels following iron depletion in HeLa cells (Fig 5E). NCOA4 levels were higher at baseline in SETD2 depleted cells and increased over time compared to the wildtype cells, consistent with perturbed ferritinophagy. FTH1 levels were relatively preserved following SETD2 depletion, and the increased ratio of NCOA4/FTH1 levels in SETD2 depleted cells was also consistent with SETD2 loss impairing ferritinophagy (Fig. 5E).

To further delineate the involvement of SETD2 in ferritinophagy, we selectively depleted autophagy adaptors and examined if they abrogated the effect on SETD2 on NCOA4. Knockout of ATG9A, an autophagy component required for ferritinophagy (*59*), increased NCOA4 levels as expected (**Fig. 5F**). SETD2 depletion in ATG9 null cells did not alter NCOA4 further (**Fig. 5F**), consistent with SETD2 loss impairing ferritinophagy in an ATG9-dependent manner. However, ATG16 null cells, which have impaired phagophore expansion but not ferritinophagy (*59–61*), showed no increase in NCOA4 levels but SETD2 loss still increased NCOA4 abundance (**Fig. 5F**), consistent with SETD2 regulating ferritin flux via ferritinophagy.

Why a chromatin-associated methyltransferase involved in transcriptional activation, and fidelity and splicing should regulate ferritinophagy was not clear. Therefore, we examined if SETD2 loss altered gene transcription, the transcriptional fidelity of IRE-containing loci or splicing of genes encoding IREs using available RNA-seq and ChIP-seq data sets for SETD2 depletion (*62*). We generated a list of iron-relevant genes using gene ontology designations (**data S1**) and examined SETD2 loss altered transcription of key iron regulatory genes or ferritinophagy adaptors. H3K36me3 levels were globally following SETD2 depletion, but no specific differences were observed at genes involved in iron homeostasis (**fig. S6A**). The mRNA levels of genes involved in iron regulation were also not altered by SETD2 loss (**fig. S6B** and **C**). We next considered if SETD2 was required for the transcriptional fidelity of IREs, which reside within non-coding regions. However, no differences in mRNA level in genes with known or predicted IREs were observed in the RNA-seq datasets (**fig. S6B** and **data S1**) (*63, 64*). SETD2 loss also did not affect the transcription levels of the *TFRC* IREs (**fig. S6D**). Therefore, it was likely that SETD2 was required for correct splicing of ferritinophagy-related genes. Consistent with this notion, SETD2 loss resulted in a general reduction in splicing events, including a pronounced reduction in exon skipping (*65*), and perturbed splicing events (**Fig. 5G**). Isoform analysis of key *NCOA4* identified differential expression when SETD2 was depleted (**fig. S6E**) (*66*). These findings indicate that genes involved in ferritinophagy are susceptible to aberrant splicing when SETD2 is depleted or inhibited.

### Loss of SETD2 drives resistance to ferroptosis in cancer cells

Pharmacological or genetic disruption of iron availability can reduce activation of ferroptosis (**Fig. 6A**) (*22, 67*). Given that inducing ferroptosis is a potential therapeutic strategy for targeting tumours, and that *SETD2* is frequently mutated in cancer, we asked if SETD2 loss may render some cancers resistant to ferroptosis. We focused on ccRCC, as a cancer where *SETD2* mutations contribute to tumorigenesis (*68*), and lung adenocarcinoma, where *SETD2* mutations are also known to occur (*69*). In both these cancer types, high SETD2 expression correlates with worse patient outcomes (**Fig. 6B**) (*70*).

**Fig. 6:**
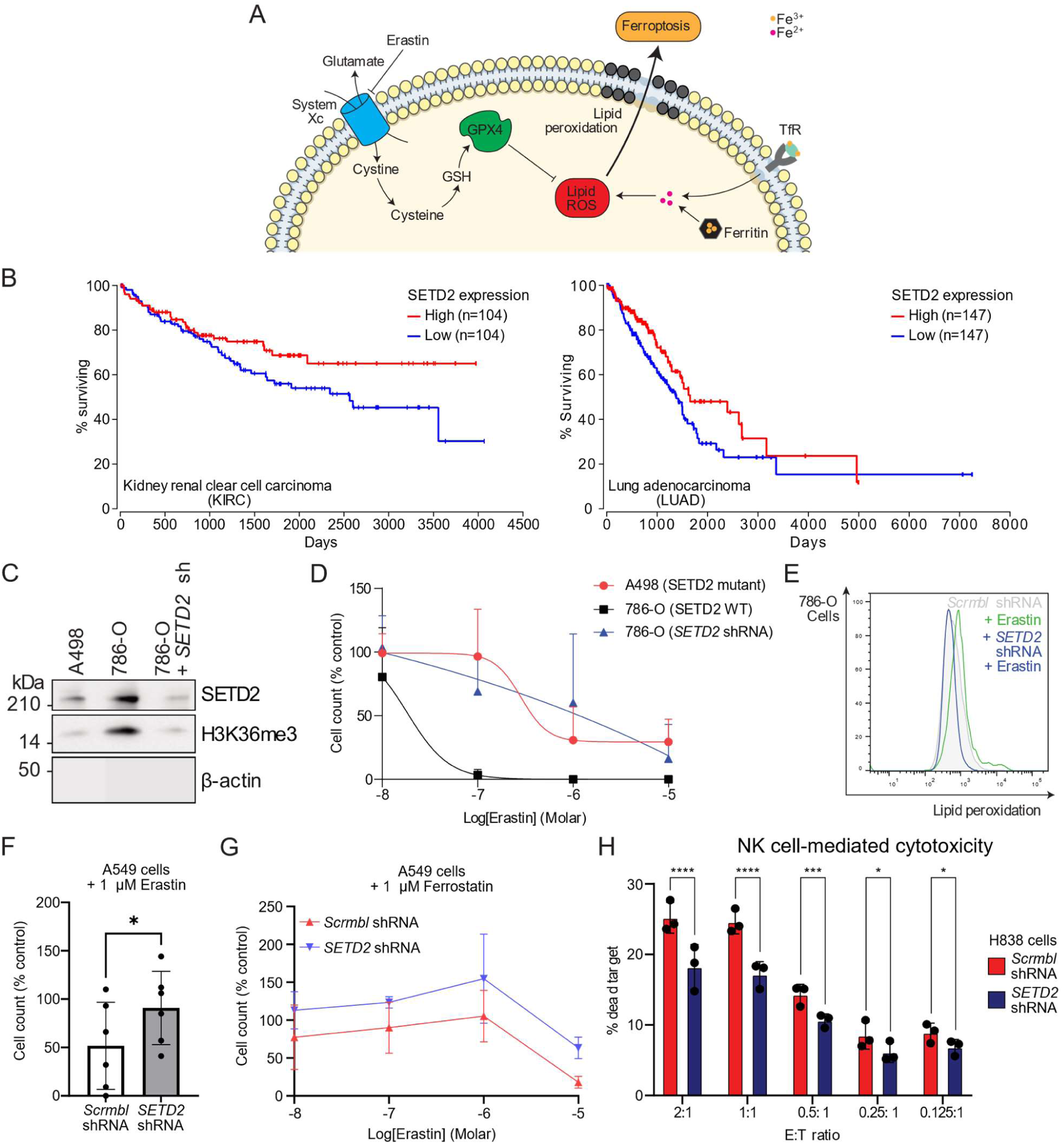
SETD2 depletion correlates with cancer cell survival and resistance to ferroptosis. (**A**) Schematic describing the key pathways in ferroptosis (membrane icon from Servier https://smart.servier.com/, CC-BY 3.0 Unported). (**B**) Kaplan-Meier survival analysis for ccRCC and lung adenocarcinoma comparing survival of patients with tumours with high (top 30%) and low (bottom 30%) *SETD2* expression levels. Curves were generated using OncoLnc with data from The Cancer Genome Atlas, logrank p-value=0.02 (KIRC) and 0.04 (LUAD). (**C**) 786-O cells underwent knockdown of *SETD2* with shRNA and were analysed by immunoblot in comparison to 786-O WT and A498 WT cells (n=3). (**D**) A498 cells and 786-O cells were treated with increasing concentrations of erastin and live cells counted after 24 hr. 786-O cells were transduced with shRNA targeting *SETD2* and the assay repeated (n=5, as percentage of untreated cells). (**E**) Control 786-O cells or those treated with *SETD2* knockdown were treated with erastin (1 μM, 24 hr) and stained with BODIPY C11 (5 μM, 35 minutes) before analysis by flow cytometry (n=3). (**F**) A549 cells with and without knockdown of *SETD2* were treated with erastin (1 μM, 48 hr) and live cells counted (n=5, as percentage of untreated cells; p=0.03, paired *t* test). (**G**) A549 cells with and without knockdown of *SETD2* were treated with increasing concentrations of erastin (10 nM-10 µM, 48 hr) and ferrostatin (1 µM, 48 hr) and live cells counted (n=3, as percentage of untreated cells). (**H**) H838 cells ± *SETD2* knockdown were incubated with primary ex vivo expanded human NK cells for 4 hr at a range of effector:target cell ratios, with both cell groups labelled (CFSE or Tag-IT Violet™). Proportion of dead target cells was measured by Fixable Viability Dye eFluorTM 780 staining and flow cytometry (n=3 biological replicates; p<0.0001, p<0.0001, p=0.0007, p=0.0147, p=0.0358, two-way repeated measures ANOVA with a post-hoc test for each E:T ratio).

We first tested if known *SETD2* mutations altered cell death by ferroptosis in ccRCC, using A498 cells, which are SETD2 deficient due to loss of one allele on the short arm of chromosome 3 (3p) and a frameshift mutation in the second allele (A498 *SETD2 -/-*), and 786-O ccRCCs which encode a functional *SETD2* (**Fig. 6C**). We treated these cells with increasing concentrations of the ferroptosis inducer, erastin, and measured cell survival after 48 hr (**Fig. 6D**). A498 *SETD2* -/- cells were resistant to erastin treatment compared to the 786-O *SETD2 +/+* cells (log IC_50_ -6.5 vs-10.4). To confirm that this difference in ferroptosis susceptibility was due to SETD2 levels, we depleted SETD2 in 786-O cells. Now, we observed a similar resistance to erastin treatment in 786-O *SETD2 -/-* as A498 *SETD2 -/-* cells (**Fig. 6D**). We also observed decreased lipid peroxidation by BODIPY-C11 staining in 786-O *SETD2 -/-* cells (log IC_50_ -5.8) (**Fig. 6E**), consistent with decreased ferroptosis. To confirm that the effects of SETD2 depletion were not just confined to ccRCC, we examined the effects of SETD2 depletion A549 lung adenocarcinoma cells. Again, SETD2 depletion resulted in resistance to ferroptosis (**Fig. 6F**), and interestingly, co-treatment with the ferroptosis inhibitor ferrostatin (1 µM) was synergistic with SETD2 depletion (**Fig. 6G**).

Inducing ferroptosis in cancers is being pursued for therapeutic effect, but ferroptosis also contributes to cancer immunosurveillance (*26*). Therefore, to explore if SETD2 may enhance cancer cell killing by immune cells, we focused on NK cells, as enhanced cancer cell killing has been reported with ferroptosis agents (*30*). We isolated NK cells from human donors and first established which cancer cells were susceptible to NK killing. Of the kidney and lung adenocarcinoma lines tested, H838 lung adenocarcinoma cells proved to be a cell type susceptible to NK killing, and this susceptibility is in keeping with prior work (*71*). We then measured NK-mediated cytotoxicity against SETD2 proficient or deficient H838 cells. SETD2 depletion reduced NK killing of H838 cells, with most marked changes observed higher effector:target ratios (**Fig. 6H**). Therefore, SETD2 depletion both enhances resistance to ferroptosis induction and may contribute to how some cancers evade immunosurveillance.

## Discussion

How iron flux is controlled in cells has been difficult to address. In part, this relates to technical limitations in measuring free iron or estimating the labile iron pool. Our generation of an endogenous IRP2 fluorescent reporter allows measurement of perturbations in iron flux, and the genetic screens using this reporter provide a comprehensive annotation of the key processes involved. Transferrin receptor uptake and recycling dominate these screens, as anticipated, but we also identified an unexpected role for mitochondrial signalling back to the cytosol, and uncovered a nuclear component in controlling iron flux, via the SETD2 histone methyltransferase.

Our data indicate that ferritinophagy is particularly susceptible to SETD2 loss, but how precisely this relates to the catalytic activity of SETD2 remains to be fully determined. Aberrant splicing may explain the uncoupling of ferritin breakdown, where NCOA4 levels accumulate along with a relative cellular iron deficiency. Why this pathway should be more susceptible than other cellular processes is unclear, but SETD2 mis-splicing has been reported for other autophagy genes unrelated to ferritinophagy (*72–74*).

Our results regarding the susceptibility to ferroptosis of ccRCC cell lines highlight why SETD2 mutations may be advantageous in the evolution of the cancer. They are also in keeping with work profiling the sensitivity of ccRCC lines to erastin-induced cell death (*75*), where A498 *SETD2 -/-* cells were more resistant to erastin than 786-O *SETD2 +/+* cells. The effect of SETD2 mutations on ferroptosis has broader disease implications, as *SETD2* mutations occur in approximately 10% of lung adenocarcinomas (*69*). However, determining the downstream consequences for *SETD2* mutations in each cancer type will be important, as death by ferroptosis will depend on other pathways involved. For example, the ISC biogenesis enzyme NFS1 is highly expressed in lung adenocarcinoma, where it protects against the induction of ferroptosis by oxidative damage resulting from high oxygen (*27*). Conversely, epidermal growth factor receptor (EGFR) mutated non-small-cell lung cancers can be particularly sensitive to the induction of ferroptosis after cystine depletion (*29*). Therefore, it will be of interest in future work to understand the relationship of SETD2 loss with other somatic mutations involved in lung adenocarcinoma development.

Our genetic screens provide a global perspective of intracellular iron metabolism, but there are some limitations. Genes identified in the screens were not consistent across all timepoints or cell types. Whilst this is not surprising, it does suggest that different cell types can have an altered reliance on ferritin uptake versus ferritin breakdown. Finally, SETD2 loss or inhibition may protect against ferroptosis by additional mechanisms to iron depletion alone, and regulation of mediators, such as cystine, glutamate and GPX4, will require consideration in future studies. However, our work provides insights into how *SETD2* mutations may contribute to cancer evolution through resistance to ferroptosis and NK cell evasion, and this will be important to consider in the context of ferroptosis-inducers as cancer treatments.

## Acknowledgments

We thank all members of the Nathan laboratory for their helpful comments on the work and manuscript. The authors gratefully acknowledge the NIHR BRC flow cytometry facility. We are grateful for the work of Anna L. Miles in the initial design of the IRP2-Clover reporter molecular constructs. We appreciate the help provided by Jason Day in the performance of ICP-MS. We also thank David Rubinsztein for the HeLa ATG9 and ATG16 null cells and for helpful discussions. This work was funded by a Wellcome Senior Clinical Research Fellowship to JAN (215477/Z/19/Z), a Wellcome Trust Clinical Training Fellowship to AWM (220535/Z/20/Z), and a Lister Institute Research Fellowship to JAN.

## Author contributions

Conceptualization: AWM, JAN Methodology: AWM, NW, JAN Investigation: AWM, CW, JJS, TV Visualization: AWM, NW Supervision: JAN, AS Writing—original draft: AWM, JAN Writing—review & editing: all authors

---

**Fig. S1.**
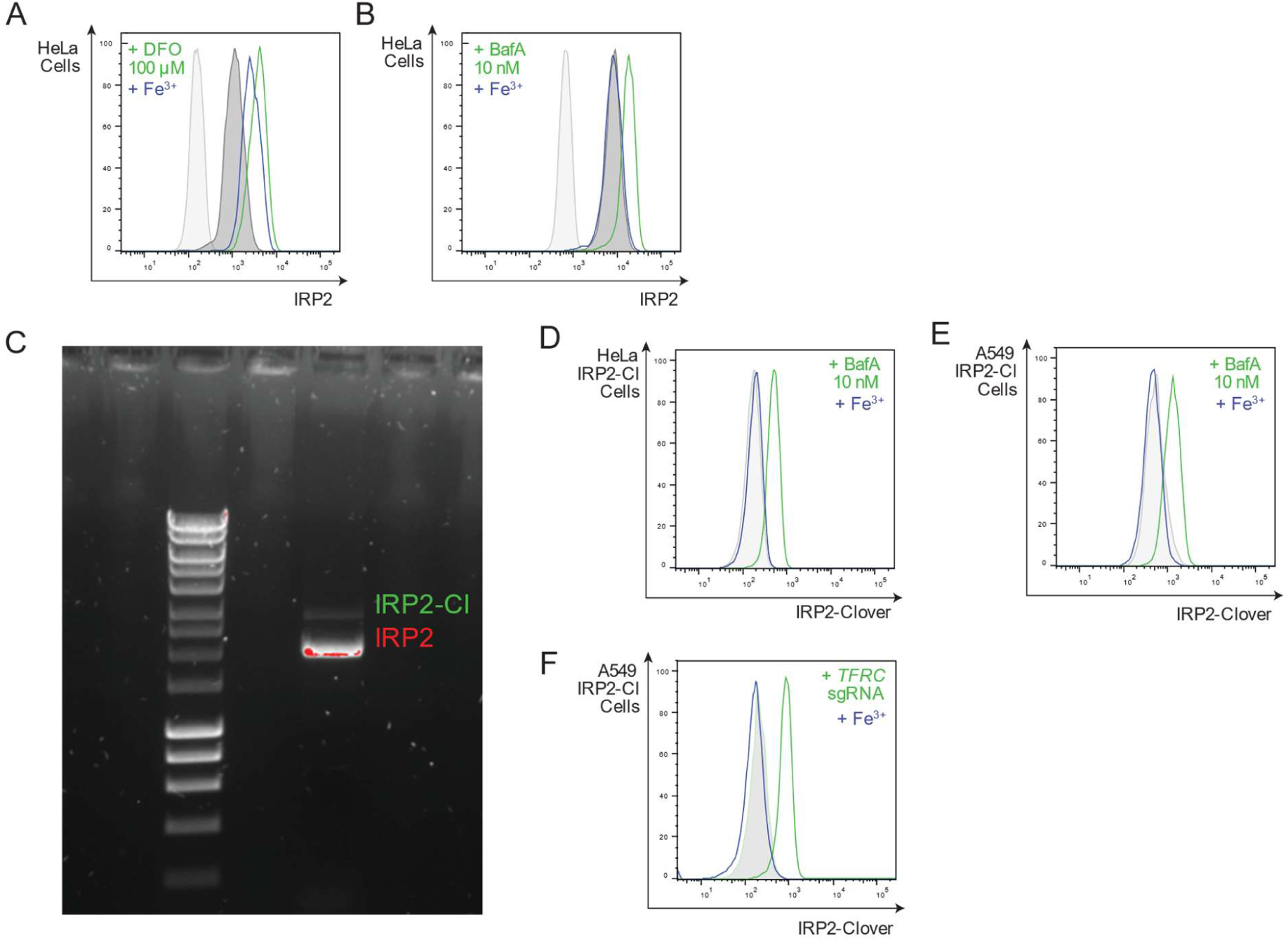
(**A**, **B**) HeLa cells were treated with DFO (**A**, 100 µM, 20 hr) or BafA (**B**, 10 nM, 20 hr) ± iron supplementation (ferric citrate, 100 µM, 20 hr), fixed and permeabilised, and stained with primary anti-IRP2 antibody (1:200) and a secondary fluorescent antibody or a secondary antibody only control (light grey), prior to analysis by flow cytometry (n=3). (**C**) DNA from clonal A549 IRP2-Clover knock-in cells was amplified via PCR. Amplicon indicating insertion of Clover (Cl) is shown (IRP2-Cl). (**D**) HeLa IRP2-Cl cells were treated with BafA (10 nM, 20 hr) ± iron (FAC, 100µM, 20 hr) and analysed by flow cytometry (representative of at least 3 independent experiments). (**E**) A549 IRP2-Cl cells were treated with BafA (10 nM, 20 hr) ± iron (FAC, 100µM, 20 hr) and analysed by flow cytometry (representative of at least 3 independent experiments). (**F**) A549 IRP2-Cl cells were transduced with sgRNA targeting *TFRC* ± iron (FAC, 100µM, 20 hr) before analysis by flow cytometry (representative of at least 3 independent experiments). *DFO=desferrioxamine, BafA=bafilomycin A, IRP2-Cl=IRP2-Clover*.

**Fig. S2.**
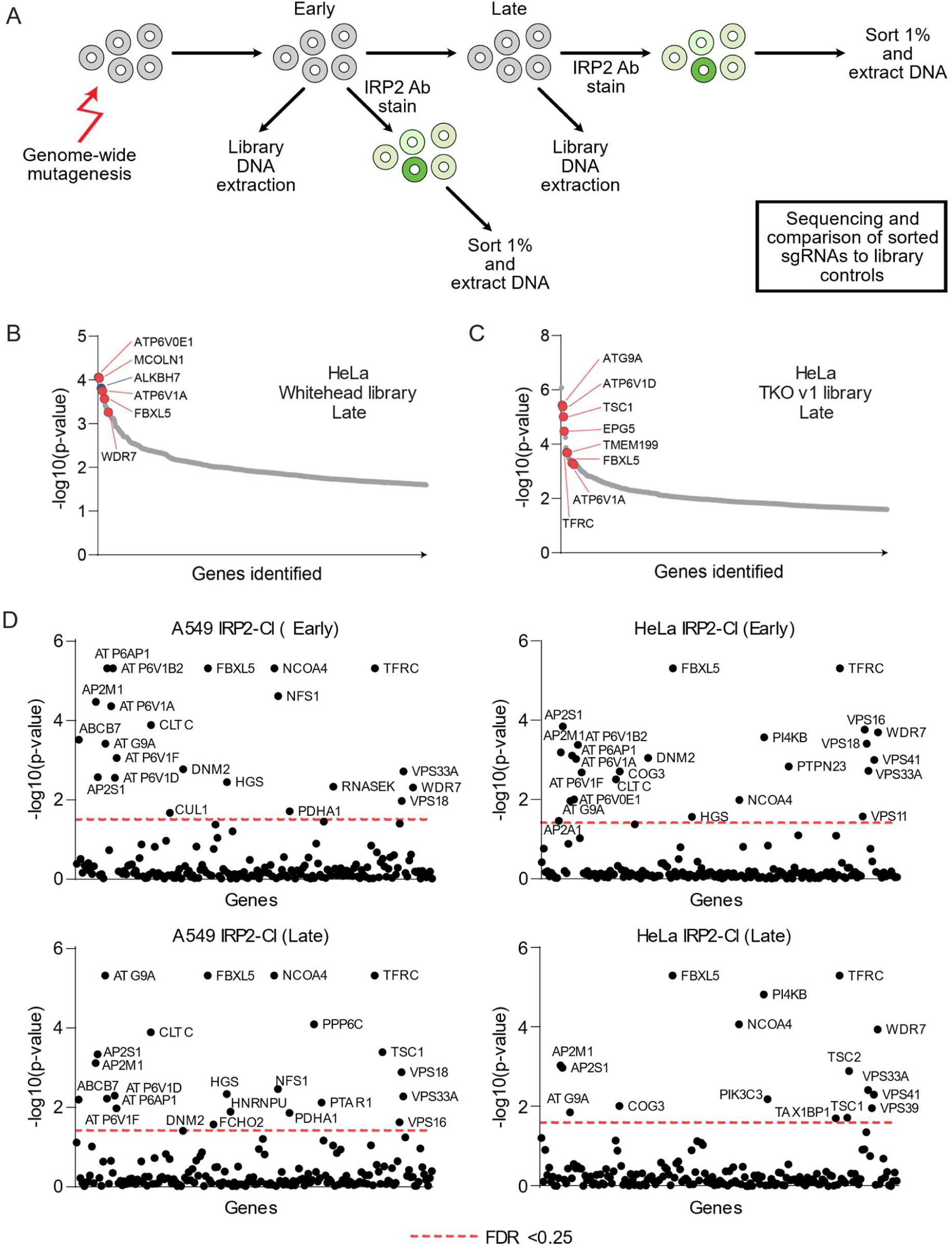
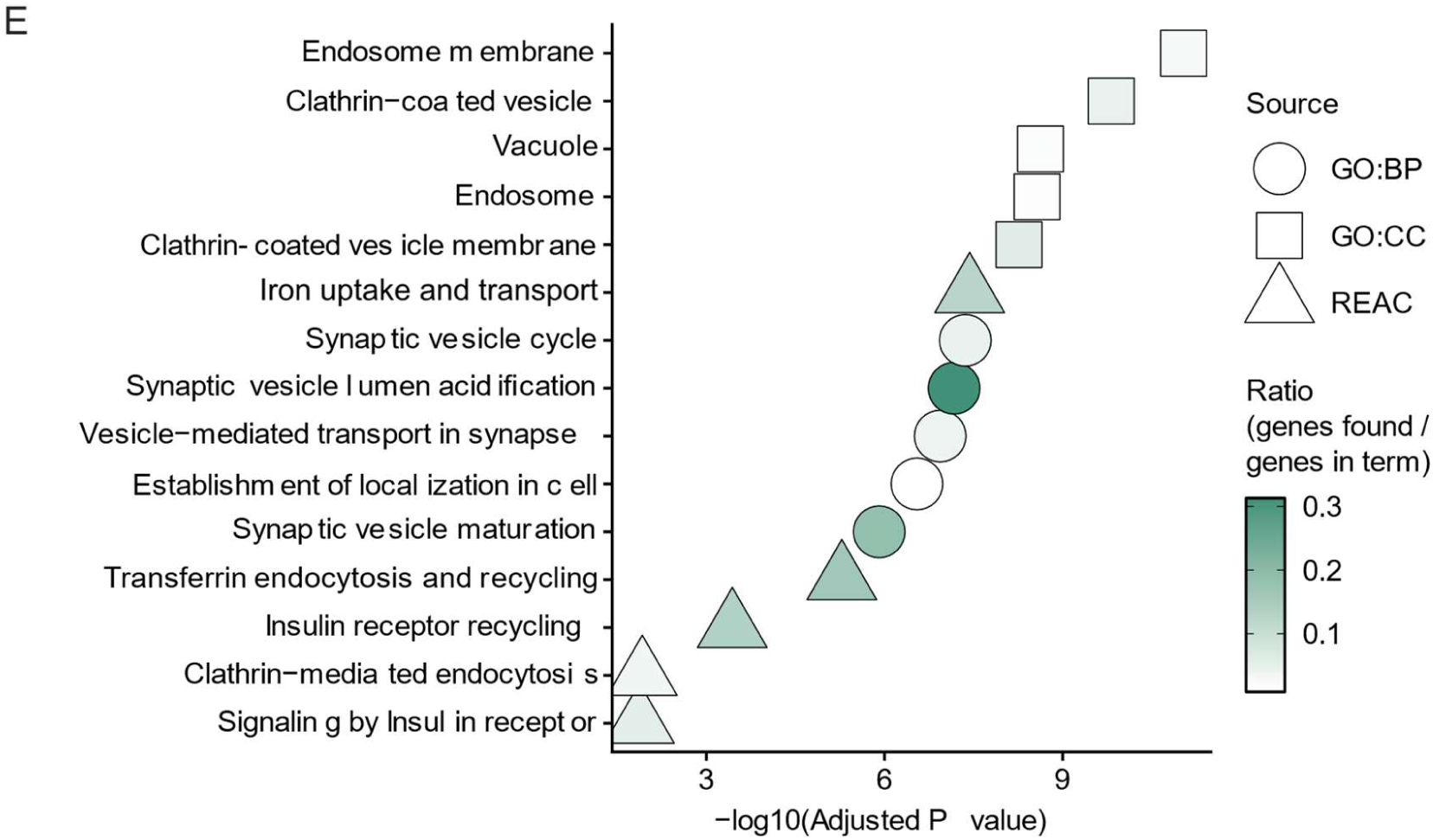
(**A**) Schematic depicting workflow for an intracellular antibody staining forward genetic screen. (**B**, **C**) Whitehead (**B**) or TKOv1 (**C**) genome-wide CRISPR libraries were transduced into HeLa cells. DNA extracted from top 1% IRP2HIGH cells by FACS was extracted and compared to library controls by MAGeCK. Bubble plots illustrate top candidate genes by -log10(p-value) at a late time point sort (day 14-15). (**D**) A549 IRP2-Cl and HeLa IRP2-Cl cells were transduced with a sub-pooled validation library composed targeting 188 candidate genes from initial mutagenesis screens. Analysis of DNA from sorted IRP2^HIGH^ cells compared to non-selected controls was undertaken using MAGeCK. Results are presented by -log10(p-value) (y axis) and gene name (x axis), with hits below a false discovery rate of 0.25 also illustrated and both early and late time points. (**E**) Gene ontology analysis of top hit candidate genes from the A549 IRP2-Clover TKOv3 screen, generated using g:Profiler. *GO:BP=Gene Ontology: Biological Process, GO:CC=Gene Ontology: Cellular Compartment, REAC=reactome*

**Fig. S3.**
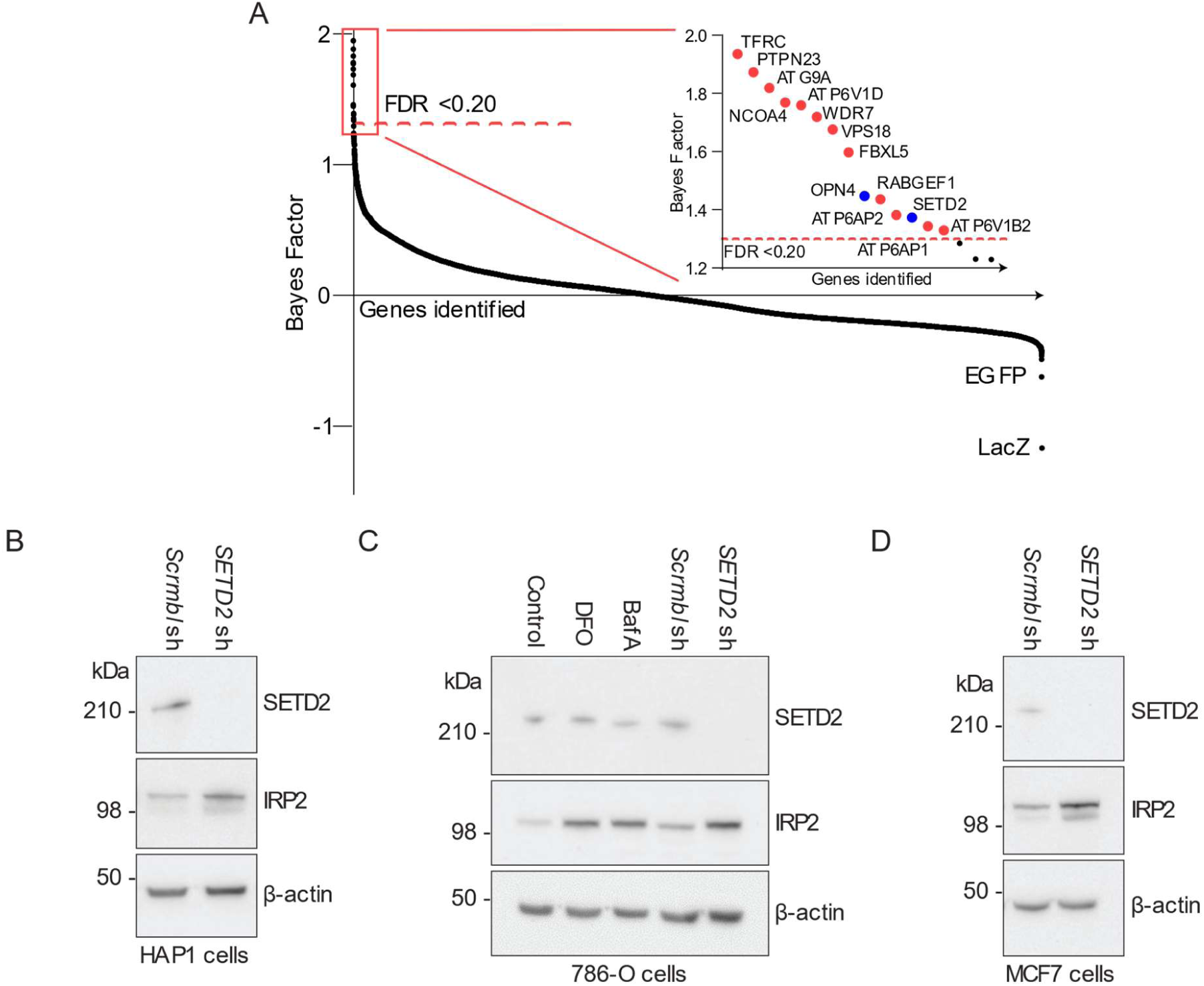
(**A**) BAGEL analysis of A549 IRP2-Clover TKOv3 CRISPR screen. DNA extracted from IRP2-Clover^HIGH^ cells at the day 16 second sort was compared to library controls. Inset magnifies top 20 genes (FDR <0.20). EGFP and LacZ are controls within the TKOv3 sgRNA library. (**B**, **C**, **D**) HAP1 (n=3) (**B**), 786-O (n=3) (**C**), and MCF7 (n=2) (**D**) cells were transduced with shRNA targeting *SETD2* or a scrambled shRNA control, and protein levels of SETD2, IRP2, and β-actin were measured by immunoblot. *DFO=desferrioxamine, BafA=bafilomycin A*.

**Fig. S4.**
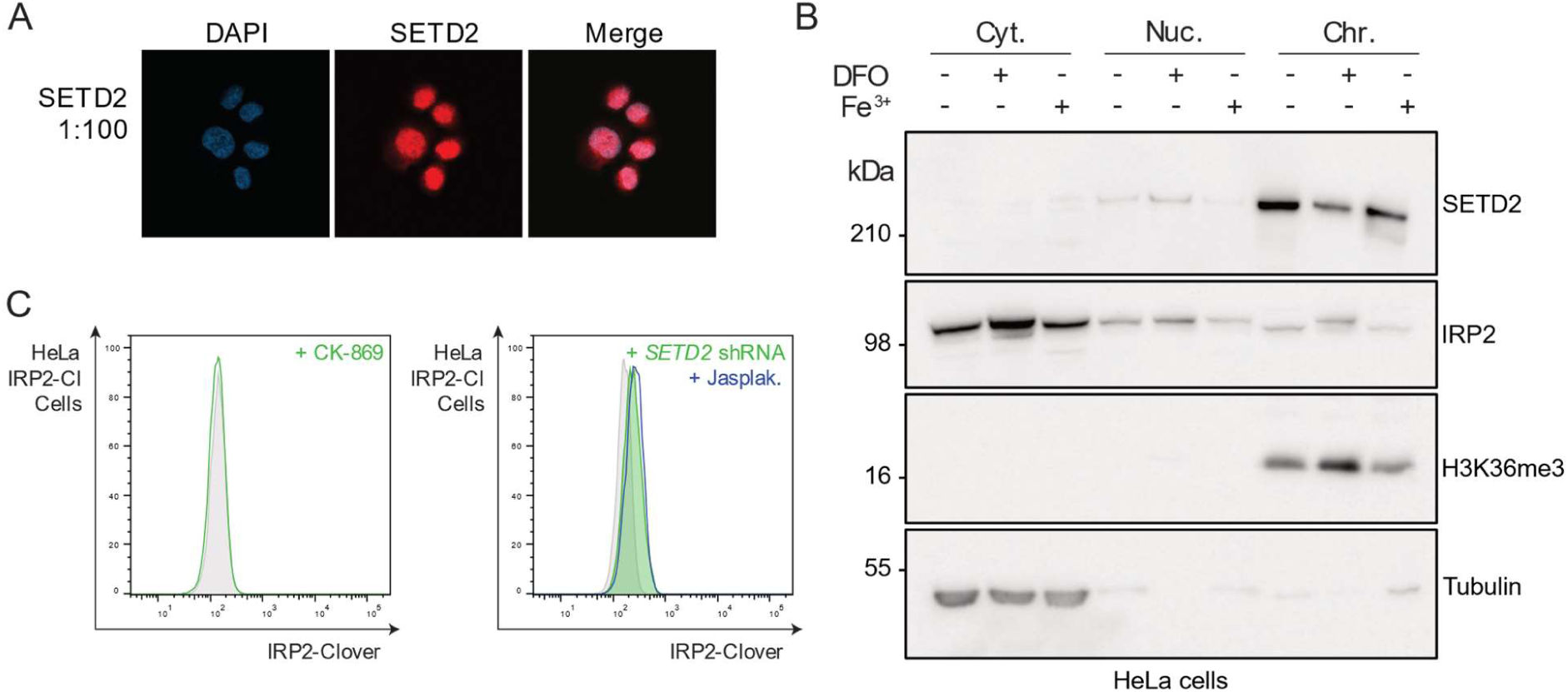
(**A**) Micrograph of HeLa cells stained with primary anti-SETD2 antibody (1:100) and DAPI (n=2). (**B**) HeLa cells were treated with iron chelation (DFO, 100 µM, 20 hr) or ferric iron excess (FAC, 200 µM, 20 hr) before subcellular fractionation. Cytosolic, nucleosolic, and chromatin fractions were analysed by immunoblot for SETD2, IRP2, H3K36me3 and Tubulin (n=3). (**C**) HeLa IRP2-Clover cells were treated with CK-869 (left panel; 200 nM, 24 hr) or transduced with shRNA targeting *SETD2* followed by treatment with Jasplakinolide (right panel; 100 nM, 24 hr) before analysis by flow cytometry (n=3). *IRP2-Cl=IRP2-Clover*.

**Fig. S5.**
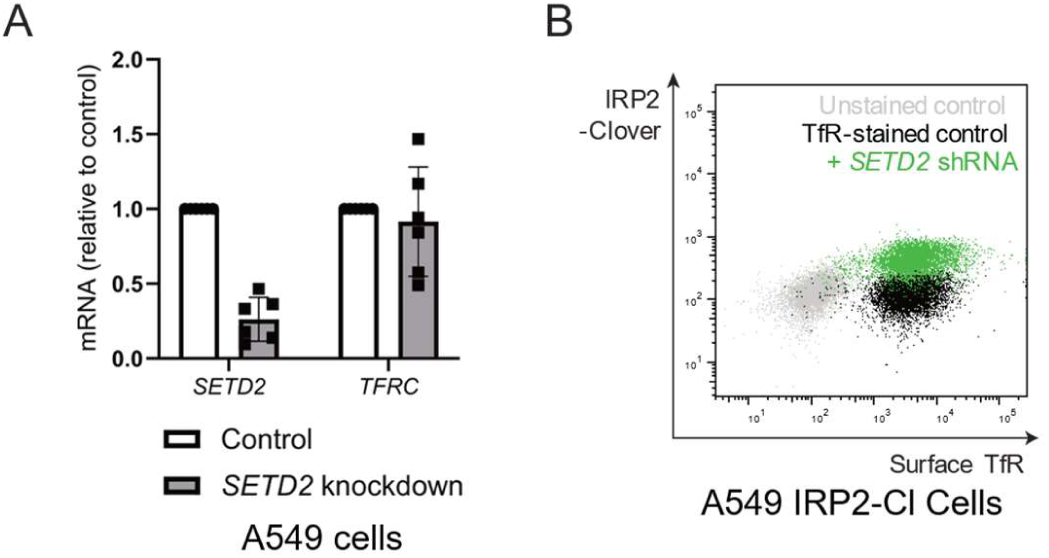
(**A**) A549 IRP2-Clover cells were transduced with shRNA targeting *SETD2*. Transcript levels of *SETD2* and *TFRC* were measured by RT-qPCR (n=6). (**B**) Cell surface staining for TfR was performed and surface TfR and IRP2-Clover levels were measured by flow cytometry (n=3).

**Fig. S6.**
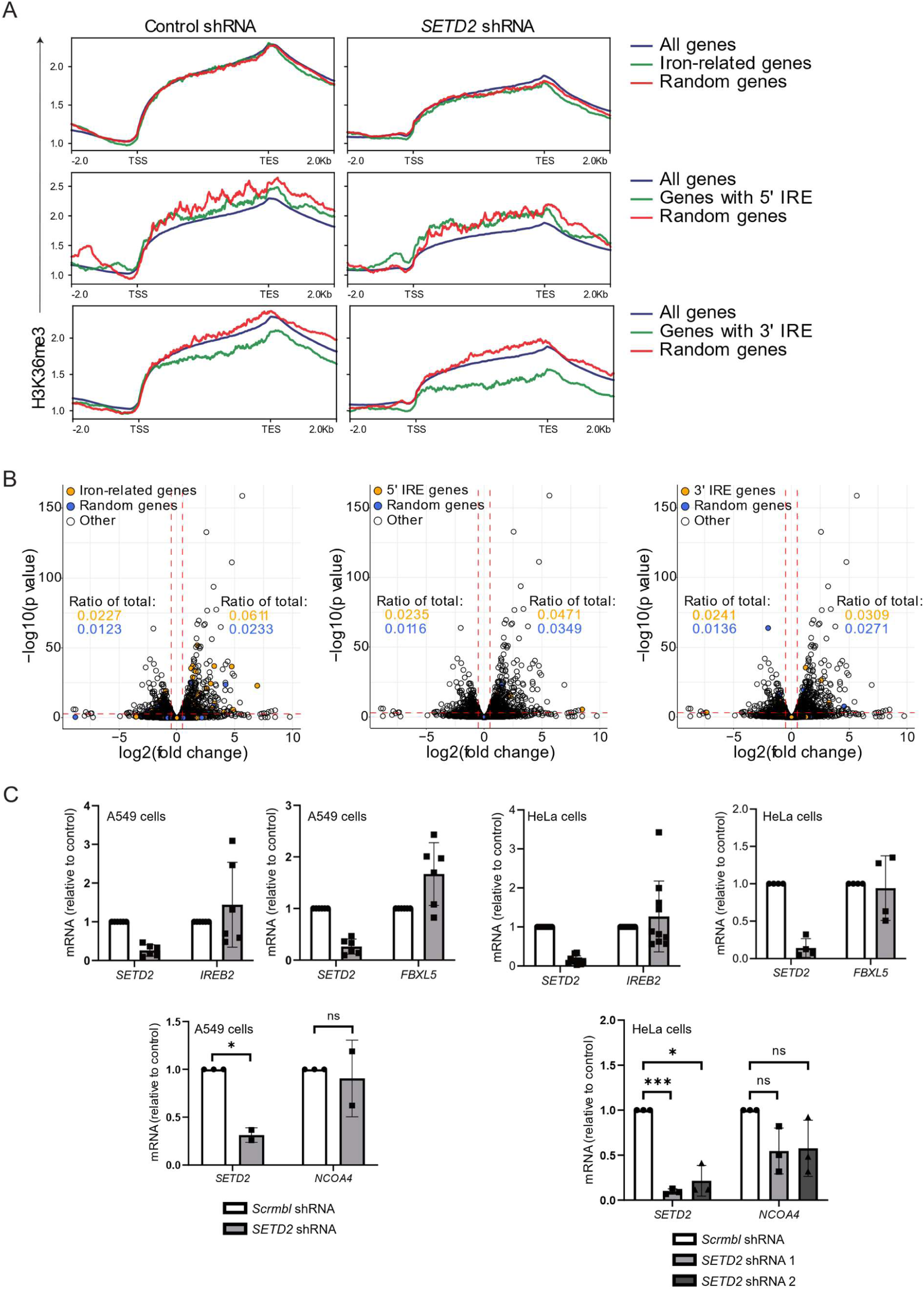
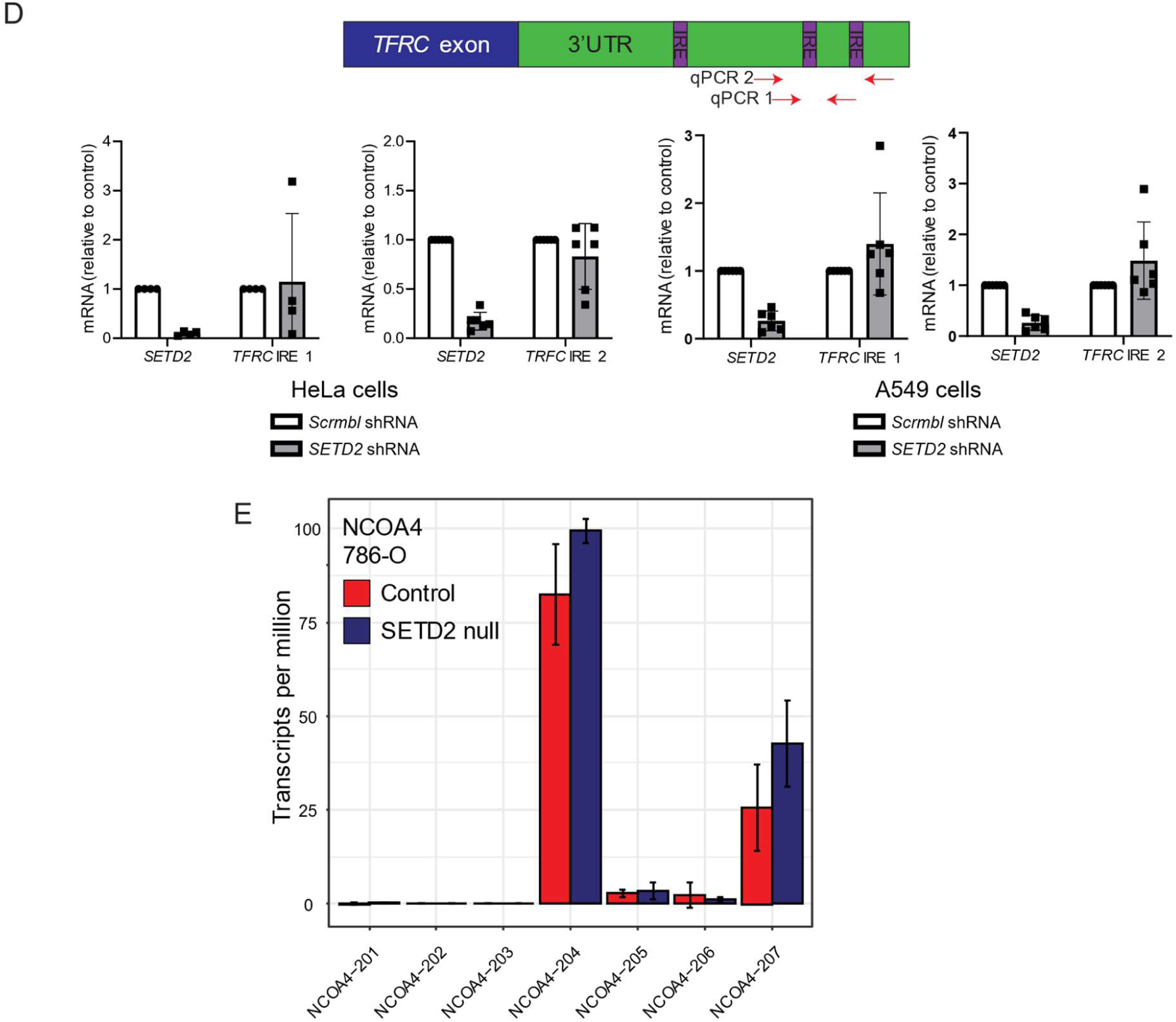
(**A**) ChIP-Seq analysis of publicly available data from HepG2 cells treated with *SETD2* or control knockdown. H3K36me3 across genes is shown for all genes, and custom datasets of iron-related genes (top panel, 731 genes), genes predicted to contain 5’ IREs (middle panel, 86 genes) and genes predicted to contain 3’ IREs (bottom panel, 295 genes). A group of randomly selected genes equal to the number of genes in each dataset served as controls. Data extracted from GSE110323. (**B**) RNA-Seq analysis of publicly available data from HepG2 cells treated with *SETD2* or control knockdown. Volcano plots showing up or down-regulated transcripts with highlighted custom datasets of iron-related genes (left panel, 731 genes), genes predicted to contain 5’ IREs (middle panel, 86 genes) and genes predicted to contain 3’ IREs (right panel, 295 genes). A group of randomly selected genes equal to the number of genes in each dataset served as controls. Data extracted from GSE110323. (**C**) A549 or HeLa cells with transduced with shRNA targeting *SETD2* or a scrambled control, and RT-qPCR was undertaken to assess transcript levels of *SETD2, IREB2* (n=6 and n=10), *FBXL5* (n=6 and n=4) and *NCOA4* (n=2 and n=3). (**D**) Schematic showing design of primers targeting IREs within *TFRC*. HeLa and A549 cells were transduced with shRNA targeting *SETD2* or a scramble control and RT-qPCR undertaken to measure levels of *SETD2* and both IRE-adjacent regions of *TFRC* transcript. (**E**) RNA-Seq analysis of publicly available data from wild type and SETD2 stable knockout (SETD2 null) 786-O cells. Transcript isoform abundance of *NCOA4* was analysed using Salmon (p=0.0459 for column factor, two-way ANOVA). Data extracted from GSE150609.

**Fig. S7.**
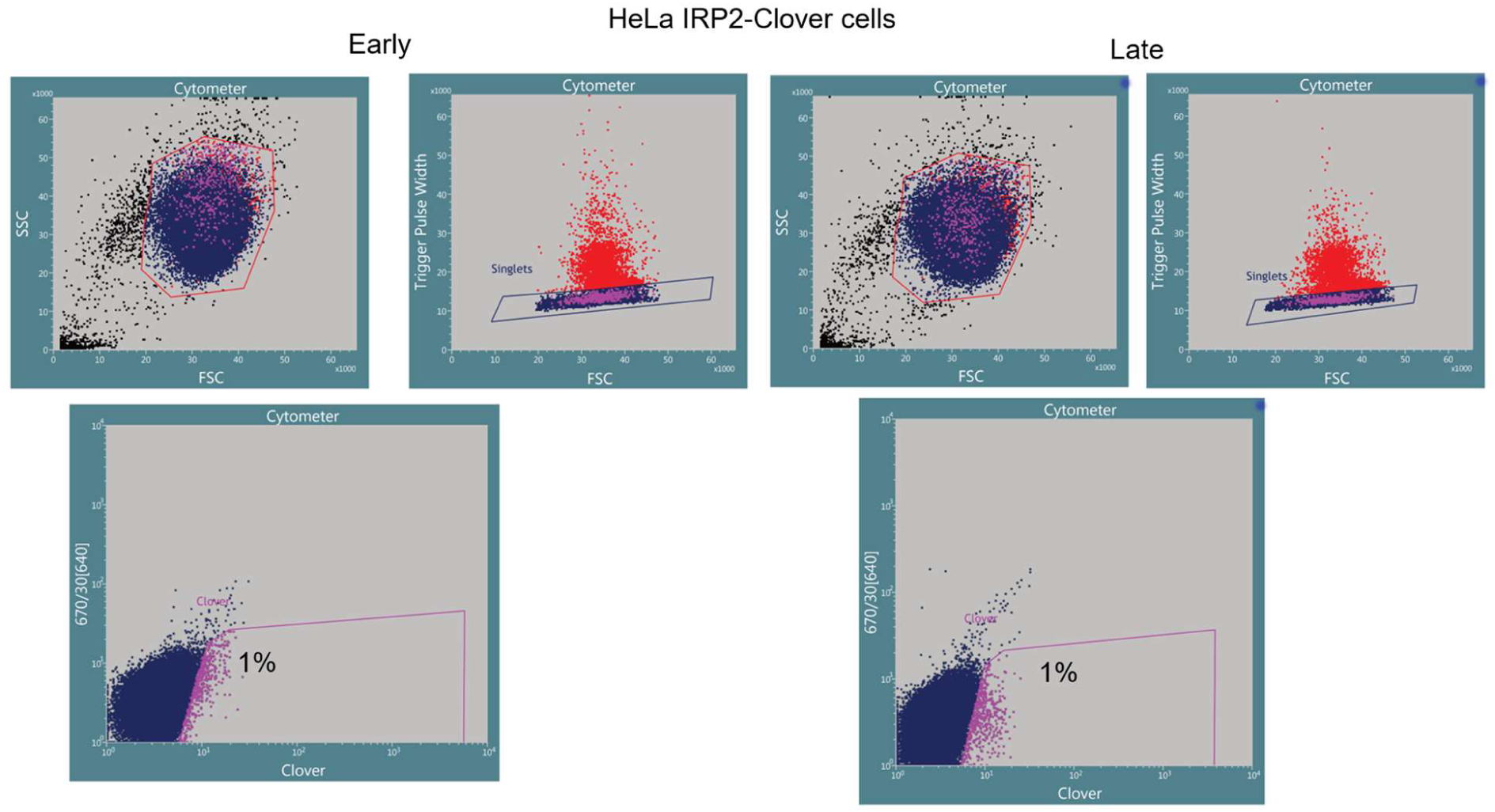
Example gating strategy for a CRISPR/Cas9 forward genetic screen. Data from HeLa IRP2-Clover screen with Whitehead library at early and late time points.

## Materials and Methods

### Cell culture

HeLa, A549, HEK293T, MCF7, HepG2, and A498 cells were maintained in Dulbecco’s modified Eagle medium (DMEM, Sigma D6429) supplemented with 10% fetal bovine serum (FBS, Sigma F0392). 786-O and H838 cells were maintained in RPMI (Sigma R8758) supplemented with 10% FBS. HAP1 cells were maintained in Iscove’s modified Dulbecco’s medium (IMDM, Gibco 31980030) supplemented with 10% FBS. Iron levels in media supplemented with FBS are approximately 5 µM (*76*). Culture media was not supplemented with Penicillin-Streptomycin outside of cell sort recovery. All cells were maintained in a 5% CO2 incubator at 37°C. HeLa ATG9 null, HeLa ATG16 null and their paired control cells were gifts from David Rubinsztein (CIMR, Cambridge). A498 cells were a gift from Peter Schraml (Zurich). Mycoplasma testing was performed on a monthly basis (MycoAlert, Lonza LT07-318). All cells were authenticated (Eurofins).

### Immunoblotting

Cells were lysed in an SDS lysis buffer (2% SDS, 50 mM Tris pH 7.4, 150 mM NaCl, 10% glycerol and 1:200 DENARASE) on ice. Electrophoresis and transfer were performed before incubation with primary and secondary antibodies (**table S1**). Immunoblots were developed using Pierce ECL or West Pico Plus before imaging (ThermoFisher 32209, 34577) via iBright (ThermoFisher) or ChemiDoc MP Imaging System (BioRad 17001402). Immunoblots are representative of at least 3 independent experiments unless otherwise noted.

### Subcellular fractionation

Cells were lysed in Buffer A (10 mM HEPES, 1.5 mM MgCl_2_, 10 mM KCl, 0.5 mM DTT and EDTA-free protease cocktail tablet, Roche) with 0.1% NP40 and incubated (10 min, shaking, RTP). 50 μl of was taken as a total cell lysate control and the remainder underwent centrifugation (1400G, 4 min, 4°C), followed by collection of the supernatant (cytosolic fraction). The nuclear pellet was resuspended in Buffer B (20 mM HEPES, 1.5 mM MgCl_2_, 300 mM NaCl, 0.5 mM DTT, 25% glycerol, 0.2 mM EDTA, EDTA-free protease inhibitor cocktail tablet) (10 min, on ice), and centrifuged (1700G, 4 min, 4°C). The supernatant (nucleosolic fraction) was collected and the remaining pellet was lysed in SDS with DENARASE® (chromatin fraction).

### Immunofluorescence

Cells were seeded on FBS-coated coverslips in 24-well plates. After 24 hr, cells were washed (Dulbecco’s PBS, DPBS, Merck D8662), fixed in 4% paraformaldehyde, permeabilised with 0.1% Triton (10 min), blocked in 4% FBS in DPBS (1 hr, RTP) and incubated with primary antibody in 40 μl blocking buffer (overnight, 4°C) (**table S1**). Coverslips were returned to the plate and washed before incubation with secondary antibody in 200 μl of blocking buffer for (1.5 hr, RTP). Coverslips were mounted on slides with a glycerol-based mount containing DAPI (CitiFluor™ AF1 + DAPI, Electron Microscopy Sciences D17970) and kept at 4°C until analysis (Zeiss LSM 980). Images were processed using ImageJ.

### Flow cytometry

Cells were resuspended in PBS prior to flow cytometry (BD Fortessa; software: FACSDiva 8.0). Flow cytometry plots are representative of at least 3 independent experiments unless otherwise noted. For intracellular flow cytometry, cells were washed with PBS and fixed in 90% ice-cold methanol for 30 min. Primary antibody was then added to these fixed, permeabilised cells for 45 min (**table S1**). Secondary fluorescent antibody was added and incubated in darkness at room temperature for 30 min (**table S1**).

### Transferrin receptor recycling assay

Cells were serum-starved for 45 min, incubated with 5 µg/ml TF-AF647 for 5 min, washed with PBS and recovered in serum free medium at indicated time points. Internalized TF-AF647 and surface TfR levels were assessed by flow cytometry.

### Molecular biology

The lentiviral sgRNA expression vector pKLVU6gRNA(BbsI)-PGKpuro2ABFP was a gift from Kosuke Yusa (Addgene #50946) (*77*). The lentiviral shRNA expression vector pc-SIREN was a gift from Paul Lehner (CITIID). PCR amplification was typically performed in 50 μl reactions using Phusion® High Fidelity DNA polymerase (NEB, M0530) per the manufacturer’s protocol. Restriction digests were performed using the relevant enzymes per manufacturer instructions, typically FastDigest enzymes (ThermoFisher). DNA was purified by agarose gel electrophoresis (supplemented with SYBR Safe, ThermoFisher) and bands extracted and recovered using a Zymoclean Gel DNA Recovery kit per the manufacturer’s instructions (Zymo Research). Bacterial transformation was performed by adding plasmid to competent E. coli on ice for 20 min, before heat shock at 42°C for 45 s, followed by recovery in super optimal broth and spreading on LB agar plates containing the relevant antibiotic and overnight incubation (37°C). Individual colonies were selected and further incubated at 37°C in LB supplemented with antibiotics before plasmids were purified using either the QIAGEN QIAprep Spin Miniprep Kit or Maxi Kit per the manufacturer’s instructions. Successful cloning was confirmed by commercial sequencing (GENEWIZ, Azenta Life Sciences).

Knock-in reporter cell lines were generated by constructing a pDonor-IRP2-Clover plasmid comprising IRP2 5’ and 3’ homology arms, a Clover-Puromycin resistance vector fragment and a kanamycin resistance vector fragment with Gibson assembly primers listed in **supplementary table 2**. This construct was transfected into cells with alongside sgRNAs targeting the C-terminus or 3’-UTR of IRP2 in a transient Cas9 expression vector. Cells were selected by puromycin and then the puromycin resistance cassette was removed by treatment with Cre recombinase or a self-excising retroviral vector encoding Cre recombinase (pHR-MMPCreGFP, a gift from Jos Jonkers, Universiteit Leiden) (*78*). Single cell clones were then isolated and IRP2-Clover integration was confirmed by PCR sequencing.

### Lentiviral production and transduction

HEK293T cells were transfected at ∼70% confluence using Fugene 6 (Promega UK E2692) at a ratio of 6 μl transfection reagent to 2 μg DNA. DNA was comprised of pCMV-dR8.91 (gag/pol), the relevant expression plasmid and pMD.G (VSVG) in a 2:3:4 ratio. The supernatant was collected at 48 hours through a 0.45 μm filter. Transduction was performed by addition of a minimum of 200 μl of supernatant to cells in media, subsequent incubation at 37°C, and addition of further medium after three hours. Cells were selected after a minimum of 27 hrs with antibiotic selection applied for 48 hr. Lentiviral packaging vectors pCMV-dR8.91 (gag/pol) and pMD.G (VSVG) were gifts from Paul Lehner (CITIID).

### Knockdown

Sequences for shRNAs were generated via the Broad Institute’s RNA interference Platform library. Oligos were designed with a TTCAAGAGA hairpin sequence, purchased commercially and cloned into a pc-SIREN backbone. Plasmids were packaged into lentivirus and transduced into cells prior to assay.

Knockdown by siRNA was adapted from a protocol by Uhrigshardt et al. (2010) with consecutive transfections 24 and 48 hr prior to assay (*79*). Sequences were generated by Dharmacon (*MFRN1*) and Uhrigshardt et al. (*HSCB*) and purchased from Eurofins (scrambled control and *MFRN1*) and IDT (*HSCB*) (**table S3**).

### RT-qPCR

Total RNA was extracted using the PureLink™ RNA Mini Kit (ThermoFisher 12183018A) according to the manufacturer’s instructions. RNA was reverse transcribed using ProtoScript II Reverse Transcriptase (NEB) according to their standard protocol (NEB #M0368). Template cDNA (40 ng) was amplified using SYBR® Green PCR Master Mix (ThermoFisher 4309155) and QuantStudio™ 7 Real-Time PCR System (ThermoFisher). Transcript levels of genes were normalized to β-actin. Primer sequences were generated by PrimerBank or obtained from published protocols (**table S4**) (*80, 81*).

### CRISPR/Cas9 knockout

For stable expression of Cas9, cells were transduced with pHRSIN-FLAG-NLS-CAS9-NLS-pGK-Hygro (a gift from Richard Timms and Paul Lehner) or Lenti Cas9-T2A-Blast (a gift from Jason Moffat, Addgene #73310) (*39*). Gene-specific sgRNA sequences (**table S5**) were generated from genome-wide CRISPR/Cas9 libraries, eCRISP or Vienna BioCentre (*82, 83*) and cloned into a pKLV-U6sgRNApGKPuro-2A-BFP backbone.

### CRISPR/Cas9 forward genetic screens

The Human Two Plasmid Activity-Optimized CRISPR “Whitehead” Knockout Library was a gift from David Sabatini and Eric Lander (Addgene #1000000095) (*38*). The Toronto human knockout pooled library (TKO) was a gift from Jason Moffat (Addgene #1000000069) (*39*). The Toronto human knockout pooled library (TKOv3) was a gift from Jason Moffat (Addgene #125517) (*40*). These libraries were packaged into lentivirus and transduced into cells expressing Cas9 at MOI 0.3 with >150-fold guide coverage. Cells were pooled prior to any selection event.

For IRP2-Clover cell screens, cells were washed in PBS supplemented with HEPES 10 mM, passed through CellTrics 50 μm filters, and suspended in sort media (PBS with 10 mM HEPES and 2% FBS) on sort days, with the top 1% of fluorescent cells underwent Fluorescent-Activated Cell Sorting (FACS, BD Influx) into collection media (49% DMEM, 49% FBS, 2% Penicillin-Streptomycin). Sorted cells were split for DNA extraction (Puregene Core Kit A, Qiagen 158388) or expansion in recovery media (DMEM with 20% FBS and 1% Penicillin-Streptomycin) and then DMEM with 10% FBS, until a second sort of the enriched population, again selecting the top 1% of cells (**Fig. 2A**). An example gating strategy is shown in **supplementary figure 7**. DNA was extracted from phenotypically non-selected library controls at each time point.

For intracellular antibody staining screens, cells were seeded 24 hr prior to sorting, with fixing and methanol permeabilisation on sort days. Cells were stained with primary rabbit anti-IRP2 antibody at 1:200 and secondary AF 488 anti-rabbit at 1:1000. Cells were then filtered before FACS (BD Influx) for the top 1% of fluorescent cells, which were collected and from which DNA was extracted (QIAmp DNA Mini Kit, Qiagen 51304) (**fig. S2A**). A late time point sort was performed using cells which had not undergone FACS, again selecting the top 1% of fluorescent cells.

A two-stage PCR was completed to amplify inserts (**table S6**). Following PCR, DNA was purified (AMPure XP, Agencourt A63880), quantified via Bioanalyzer (Agilent DNA 1000, 5067-1504) and sequencing performed by Illumina MiniSeq, HiSeq or NovaSeq with a custom primer (**table S10**). DNA was extracted from phenotypically non-selected library controls at each time point.

A custom iron sublibrary containing 184 genes was developed with four top-ranking sgRNAs for each screen-identified gene generated using the Vienna Bioactivity CRISPR score (www.vbc-score.org) (*83*) (**data S1**). Chr10promiscuous, Luciferase, intergenic and non-targeting control sgRNA sequences were taken from the TKO and Whitehead libraries (*38, 39*). The sublibrary was purchased commercially (IDT) and amplified by PCR, with clean-up performed using the QIAquick Nucleotide Removal Kit (Qiagen 28306) per the manufacturer’s instructions. Guides were cloned into a pKLV backbone and transformed by electroporation into Stbl4 cells. After recovery, bacteria were serially diluted and plated overnight or continued in LB, both at 37°C. Colonies were counted to ensure representation and DNA extracted using a Maxi Kit before amplification in a two-stage PCR and sequencing via MiniSeq to confirm guide representation.

### CRISPR/Cas9 screen analysis

A Snakemake pipeline generated and curated by Niek Wit (https://github.com/niekwit/crispr-screens and https://doi.org/10.5281/zenodo.10286661) was utilized for bioinformatic analysis. Initially, the quality of the raw data was assessed using FastQC v0.12.1 and MultiQC v1.23. Reads were then quality trimmed to 20 bp using Cutadapt v4.0 (*84*). The trimmed reads were aligned to all sgRNA sequences using HISAT2. MAGeCK and BAGEL2 were employed to determine the changes in normalized read counts between samples (*85, 86*). This analysis compares DNA extracted after sorting to an unsorted DNA library. Outputs from MAGeCK and BAGEL2 are available in **supplementary data 2**.

### ChIP-Seq

A Snakemake pipeline generated and curated by Niek Wit (https://github.com/niekwit/chip-seq and https://doi.org/10.5281/zenodo.13801526) was utilized for bioinformatic analysis. After initial QC of the raw data (GSE110323 (*62*)) as above, reads were trimmed using Trim Galore v0.6.10. Bowtie2 v2.5.0 was used to align the reads to the human genome (hg38). From these BAM files, blacklisted regions were removed using BEDTools v2.31.1 (https://doi.org/10.1093/bioinformatics/btq033). BAM files were then sorted and duplicates removed using SAMtools v1.20 and Picard v3.2.0 (https://broadinstitute.github.io/picard/), respectively. To generate profile plots, deepTools v3.5.2 (https://doi.org/10.1093/nar/gkw257) subcommand plotProfile was used with BigWig files generated by the subcommand bamCoverage with BAM files as input.

### RNA-Seq

Data analysis of publicly available RNA-Seq data (GSE150609 and GSE110323) (*62, 65*) was conducted using a Snakemake pipeline (https://doi.org/10.5281/zenodo.10139567). After assessing the quality of the raw data with FastQC and MultiQC, sequence reads were trimmed to remove adaptor sequences and low-quality nucleotides using TrimGalore. Transcript quantification was carried out with Salmon against the Gencode Homo sapiens reference transcriptome/genome (build 44) (*66*). Differential transcript analysis was performed using DESeq2, identifying genes with adjusted P values <0.01 and log2(fold change) >0.5 as differentially expressed for each comparison (*87*). Lists of genes predicted to contain 5’ or 3’ IREs were obtained from Hin et al. (2021), generated using the Searching for IREs tool (SIRES) (**data S1**) (*63, 64*).

### Alternative splicing

Data analysis of publicly available RNA-Seq data (GSE150609) (*65*). Trim Galore was used to remove adapter sequences. Read alignment was performed against the human genome (hg38) with STAR (*88*). To quantify alternative splicing events, rMATS was used (*89*). A custom R script using the package Tidyverse (https://doi.org/10.21105/joss.01686) was used for plotting.

### Gene ontology

Gene ontology analysis of screen hits was undertaken using g:Profiler (*41*).

### Inductively coupled plasma-mass spectrometry (ICP-MS)

ICP-MS was developed from a protocol by Stangherlin et al. (2021) and performed in collaboration with Jason Day (Department of Earth Sciences, University of Cambridge) (*90*). For each biological replicate, approximately 5x10^5^ cells (corresponding to ∼10 ng of iron) in technical triplicate were washed in PBS before lysis (30 minutes, RTP) with high purity 65% nitric acid (Merck Millipore 100441) supplemented with 0.1 mg/L (100 ppb) cerium (Romil E3CE#) as a procedural internal standard. Prior to analysis, samples were diluted to a final concentration of 5% HNO3. Data were collected and analysed using Syngistix version 1.1, normalized based on cerium abundance and cell number (counted from duplicate cultures at time of harvest) and analysed using GraphPad prism.

### Ferroptosis assays

For manual cell counts, cells were seeded in 6-well plates, treated with erastin and counted using Trypan blue after 48 hr using a haemocytometer. To assess lipid peroxidation, cells were seeded in 6-well plates and treated with erastin 24 hr prior to staining with 5 μM BODIPY™ 581/591 C11 (ThermoFisher, D3861) in for 35 min, before resuspension in 500 μl Hank’s Balanced Salt Solution (Sigma-Aldrich H9394) and analysis by flow cytometry using a 488 nm laser with a 530/30 nm bandpass filter (BD LSR II) and data processed using FlowJo (BD Life Sciences).

### NK competitive cytotoxicity assays

Primary human NK cells were isolated from leukocyte cones obtained from anonymous donors via NHS Blood and Transplant (REC 22/PR/1280 approved by London - Chelsea Research Ethics Committee) via negative selection using the NK Cell Isolation Kit (Miltenyi Biotec) and expanded for 14 days in NK MACS medium (Miltenyi Biotec) supplemented with 5% heat inactivated human AB serum (Sigma-Aldrich), 50µg/ml Gentamicin Sulphate (ThermoFisher Scientific), 50µM 2-Mercaptoethanol (Gibco), recombinant human IL-2 (500 U/ml, Peprotech) and IL-15 (10 U/ml, Peprotech). Assays were performed in PBMC washing media (RPMI-1640 (Sigma-Aldrich), supplemented with 10% heat inactivated FBS (Sigma-Aldrich), 50µg/ml Gentamicin Sulphate (ThermoFisher Scientific) and 2mM Glutamine (Sigma-Aldrich).

Target cells were washed with D-PBS and labelled with either CFSE (0.5 nM, Invitrogen) or Tag-IT Violet^TM^ Proliferation and Cell Tracking Dye (1.25 µM, BioLegend) for 20 min at 37°C. The reaction was quenched by the addition of 5 ml PBMC washing media and incubated for 5 min at room temperature. After washing, the cells were counted and adjusted to the required density. The competitive cytotoxicity assay was set up in 96 well U-bottom plates with a total of 5x10^4^ target cells per well at a 1:1 ratio (control:SETD2sh) and a range of effector:target ratios as indicated in the figures. The plate was incubated at 37°C 5% CO_2_ for 4 h. At the end of the assay, cells were washed with D-PBS and stained with Fixable Viability Dye eFluor^TM^ 780 (ThermoFisher Scientific, 1:2000 dilution) for 5 min at 37°C. After washing with FACS buffer, cells were fixed with 2% PFA for 5 min at room temperature and washed once with FACS buffer prior to acquisition on a flow cytometer (MACSQuant VYB). Target cells were identified by CFSE or Violet and the percentage of dead cells with the Fixable Viability Dye eFluor^TM^ 780. The specific cell death was calculated by subtracting the percentage of dead cells in the target only well.

### Statistical Analysis

GraphPad Prism was used to calculate *P* values for individual experiments, including for paired *t* test and two-way analysis of variance (ANOVA) where indicated. CRISPR screens and ChIP-Seq were analysed as described in the relevant sections above. Kaplan-Meier curve data was analysed using the logrank test, generated by OncoLnc (*70*).

## Data availability

Any additional information required to reanalyse the paper is available from the author upon request.

## Supplementary information

**Table S1.**

List of antibodies. IB=immunoblot, IF=immunofluorescence, FC=flow cytometry.

**Table S2.**

List of sequences for IRP2-Clover knock-in.

**Table S3.**

List of knockdown sequences.

**Table S4.**

List of qPCR primers. Sequences generated via PrimerBank (Wang and Seed, 2003) or previously published (van Uden et al, 2011).

**Table S5.**

List of sgRNA sequences.

**Table S6.**

List of sequences for CRISPR/Cas9 screens.

**Data S1.**

Tab 1: Genes included in sub-pooled sgRNA library. Control sgRNAs from other CRISPR libraries: chr10Promiscuous (TKO), Control (Whitehead), Intergenic (Whitehead), Luciferase (TKO).

Tab 2: List of iron-related genes, defined by GO designation.

Tab 3: List of high-quality predicted 5’ IRE genes. Generated by Hin et al. (2021) using SIREs (Campillos et al. 2010).

Tab 4: List of high quality predicted 3’ IRE genes. Generated by Hin et al. (2021) using SIREs (Campillos et al., 2010).

**Data S2.**

Tab 1: MAGeCK output for A549 IRP2-Clover TKOv3 screen (early time point).

Tab 2 MAGeCK output for A549 IRP2-Clover TKOv3 screen (late time point).

Tab 3: MAGeCK output for HeLa IRP2-Clover Whitehead screen (early time point).

Tab 4: MAGeCK output for HeLa IRP2-Clover Whitehead screen (late time point).

Tab 5: MAGeCK output for HeLa IRP2 antibody Whitehead screen (late time point).

Tab 6: MAGeCK output for HeLa IRP2 antibody TKOv1 screen (late time point).

Tab 7: MAGeCK output for A549 IRP2-Clover iron sublibrary screen (early time point).

Tab 8: MAGeCK output for A549 IRP2-Clover iron sublibrary screen (late time point).

Tab 9: MAGeCK output for HeLa IRP2-Clover iron sublibrary screen (early time point).

Tab 10: MAGeCK output for HeLa IRP2-Clover iron sublibrary screen (late time point).

Tab 11: BAGEL2 output for A549 IRP2-Clover TKOv3 screen (late time point).

