## Supplement for "Mapping the genetic landscape of iron metabolism uncovers the SETD2 methyltransferase as a modulator of iron flux"

| Antibody | Source | Identifier | Dilution |
|---|---|---|---|
| β-actin | Sigma | A228 | IB: 1:30,000 |
| CD71/TfR1 | BD Pharmigen | 555534 | |
| FTH1 | Cell Signaling | #3998 | IB: 1:1,000 |
| H3K36me3 | Cell Signaling | #9763 | IB: 1:1,000 |
| IRP2 (D6E6W) | Cell Signaling | #37135 | IB: 1:1,000 |
| NCOA4 (ARA70) | Bethyl | A302-272A | IB: 1:1,000 |
| SETD2 | ABclonal | A3194 | IB: 1:1,000<br>IF: 1:100 |
| HRP anti-mouse | Jackson | 115-035-146 | IB: 1:20,000 |
| HRP anti-rabbit | Jackson | 115-035-045 | IB: 1:20,000 |
| Alexa Fluor 488 (anti-rabbit) | Invitrogen | A11034 | FC: 1:1,000<br>IF: 1:400 |
| Alexa Fluor 568 (anti-rabbit) | Invitrogen | A11036 | IF: 1:400 |
| Alexa Fluor 647 (anti-rabbit) | Invitrogen | A21245 | IF: 1:400 |
| Alexa Fluor 647 (anti-mouse) | Invitrogen | A11036 | FC: 1:1000 |

**Table S1.** List of antibodies. *IB=immunoblot, IF=immunofluorescence, FC=flow cytometry.*

| Function | Sequence |
|---|---|
| Knock-in 5' fw | TTGGGCTCCCCGGGCGCGACTAGTGAATTCCTCC TGGAATTACATTGAATATACAGGTAT |
| Knock-in 5' rv | ACCAGATCCGCCACCAGATCCGCCCGATCGTGAG AATTTTCGTGCTACAAAGTTTAATAA |
| Knock-in 3' fw | AGCATACATTATACGAAGTTATTTAATTAATATCTAC TTACAATAGATACGTTTCATAAC |
| Knock-in 3' rv | TTCTTATAATCAGCATCATGATGTGGTACCGTTCAG AGTTTAATGATTGAATAACATTCT |
| *IREB2* sgRNA | ATTCCTGGGTCCAGCACAAA |
| Sequencing fw | CTCCTGGAATTACATTGAATATACAGGTAT |
| Sequencing rv | GTTCAGAGTTTAATGATTGAATAACATTCT |

**Table S2.** List of sequences for IRP2-Clover knock-in.

| Target | Sequence (5’ → 3’) |
|---|---|
| *EXOC1* (sh) | GATGAATACCAAGAGTTAAAT |
| *HSCB* (si) | CATAGAAATAATGGAAATCAA |
| *MFRN1* (si) | GGUAAUGAAUCCAGCAGAA |
| *Scrmbl* (sh) (1) | GCATAATTAATATCCGCGTGT |
| *Scrmbl* (sh) (2) | ATGGATATATAACGACCTAGT |
| *Scrmbl* (sh) (3) | ATACGATGAATAGACGACAGC |
| *Scrmbl* (si) | CAGUCGCGUUUGCGACUGG |
| *SETD2* (1) (sh) | AGTAGTGCTTCCCGTTATAAA |
| *SETD2* (2) (sh) | ACGAATTAAAGACCGCAATAA |

**Table S3.** List of knockdown sequences.

| Primer name | Sequence (5' → 3') |
|---|---|
| *ACTB* fw | CTGGGAGTGGGTGGAGGC |
| *ACTB* rv | TCAACTGGTCTCAAGTCAGTG |
| *EXOC1* fw | CCTGCTGAGCATTGTGAATGT |
| *EXOC1* rv | CACAGGGCGTTCAGTTGTCA |
| *FBXL5* fw | AGAACACTCCACAGGTATAACCC |
| *FBXL5* rv | CTGCATCGACATAACTCTTGAGG |
| *HSCB* fw | AGAGAAGCATTCGACCCTGGT |
| *HSCB* rv | AGGAATTGCCTGTCCATTTCAT |
| *IREB2* fw | TCGATGTATCTAAACTTGGCACC |
| *IREB2* rv | GCCATCACAATTTCGTACAGCAG |
| *MFRN1* fw | GATGGGGACAGCCGAGATG |
| *MFRN1* rv | ACCGGGTACATGACCGAGT |
| *NCOA4* fw | CAGCAGCTCTACTCGTTATTG G |
| *NCOA4* rv | TCTCCAGGCACACAGAGACT |
| *SETD2* fw | TGCTTCTAGTCGATTTTTGCCC |
| *SETD2* rv | AGGGTTTGGAGTATCACTTTGC |
| *TFRC* fw | ACCGGCACCATCAAGCT |
| *TFRC* rv | TGATCACGCCAGACTTTGC |
| *TFRC* IRE1 fw | TCCAGTACCTTTGTCACAATCCT |
| *TFRC* IRE1 rv | TCCGATACAGACACTGTGGT |
| *TFRC* IRE2 fw | TCCAAGGTGTAACTCTAATTCCCA |
| *TFRC* IRE2 rv | TGTTCCCGATAATTACTTACACCC |

**Table S4.** List of qPCR primers. Sequences generated via PrimerBank (Wang and Seed, 2003) or previously published (van Uden et al, 2011).

| Target | Sequence (5’ → 3’) |
|---|---|
| *EXOC1* (1) | GCATACACCAAACTTATCAG |
| *EXOC1* (2) | GGCTAACATCCAGTCAATCA |
| *NCOA4* | CAATCTCCACACCTTTGGGC |
| *SETD2* (1) | GCGGAGCTGATACTTACTCA |
| *SETD2* (2) | GGACTGTGAACGGACAACTG |
| *TFRC* | GCTCTGGAGATTGTCTGGAC |

**Table S5.** List of sgRNA sequences.

| Function | Sequence |
| --- | --- |
| Whitehead/TKOv1 outer PCR fw | AGGGCCTATTTCCCATGATTCCTT |
| Whitehead/TKOv1 outer PCR rv | TCAAAAAAGCACCGACTCGG |
| TKOv3 outer PCR fw | GAGGGCCTATTTCCCATGATTC |
| TKOv3 outer PCR rv | CAAACCCAGGGCTGCCTTGGAA |
| Inner PCR fw | AATGATACGGCGACCACCGAGATCTACACTCTCTTGTGG AAAGGACGAGGTACCG |
| Inner PCR rv | CAAGCAGAAGACGGCATACGAGATNNNNNNGTGACTGG AGTTCAGACGTGTGCTCTTCCGATCTATTTTAACTTGCTA TTTCTAGCTCTAAAAC |
| Sequencing primer | ACACTCTCTTGTGGAAAGGACGAAACACCG |

**Table S6.** List of sequences for CRISPR/Cas9 screens.
